# ER-associated protein degradation initiates by retrotranslocation from the ER quality control compartment

**DOI:** 10.64898/2026.02.04.703732

**Authors:** Chaitanya Patel, Navit Ogen-Shtern, Haddas Saad, Oscar R. Burrone, Gianluca Petris, Gerardo Z. Lederkremer

**Affiliations:** The Shmunis School of Biomedicine and Cancer Research, Cell Biology Division, George Wise Faculty of Life Sciences, Tel Aviv University, Tel Aviv 69978, Israel; Sagol School of Neuroscience, Tel Aviv University, Tel Aviv 69978, Israel; The Skin Research Institute, The Dead-Sea and Arava Science Centre, Masada 86910, Israel; Molecular Immunology Laboratory, International Center for Genetic Engineering and Biotechnology, Trieste, Italy; Genome Engineering and Biotechnology Unit, Fondazione Italiana Fegato, AREA Science Park, Trieste, Italy; Department of Medicine, University of Udine, Udine, Italy

**Keywords:** ER-associated degradation (ERAD), ER quality control compartment (ERQC), HRD1 ubiquitin ligase, p97/VCP, BAP–BirA biotinylation

## Abstract

Misfolded proteins are eliminated from the endoplasmic reticulum (ER) by ER-associated degradation (ERAD), a process requiring their exit from the ER and delivery to cytosolic proteasomes via the key retrotranslocation step. Conventional assays detect fully extracted substrates, overlooking early initiation events. Here, we use an *Escherichia coli*-derived Biotin Acceptor Peptide (BAP)-biotin ligase BirA system to detect ERAD substrates upon cytosolic exposure. Using the model ERAD substrate H2a, we show that cytosolic BirA selectively biotinylates luminal BAP tags during early cytosolic exposure, enabling detection of substrates engaging the mammalian retrotranslocation machinery independently of subsequent extraction. Combined with high-resolution expansion fluorescence microscopy, this approach reveals active initiation of ERAD at the specialized ER-derived quality control compartment (ERQC). Substrates become transiently exposed to the cytosol while remaining membrane-associated, followed by tight coupling of polyubiquitination, transmembrane segment extraction, deglycosylation and degradation. Functional perturbations indicate that mannose trimming enables initiation, whereas p97 and HRD1 support later extraction and ubiquitination.

---

Misfolded secretory and membrane proteins are removed from the endoplasmic reticulum (ER) by the ER-associated degradation (ERAD) pathway, which ensures proteostasis by targeting aberrant proteins for ubiquitination and subsequent degradation by the cytosolic proteasomes^1–4^. Dysfunction of ERAD leads to the accumulation of misfolded proteins and ER stress, contributing to a wide range of pathological conditions, including neurodegenerative diseases, metabolic disorders, cancer, and rare genetic syndromes, underscoring the importance of understanding the molecular mechanisms that govern this pathway. ^5–7^.

A central requirement of ER-associated degradation (ERAD) is the ability of luminal and membrane-embedded substrates - including misfolded secretory and membrane proteins, as well as microbial toxins and viral or other infectious agents that exploit this pathway - to exit the endoplasmic reticulum (ER)^8–14^. This process requires retrotranslocation, whereby substrates located within the ER lumen or membrane become exposed to the cytosol during their exit from the ER. Despite extensive investigation, where and how retrotranslocation is initiated in mammalian cells remains incompletely understood. Extensive genetic, biochemical, and structural studies have identified core ERAD components, including ubiquitin ligases, cytosolic ATPases, and the proteasome. However, most experimental approaches to study retrotranslocation detect substrates only after complete extraction into the cytosol.^15–18^. As a result, the earliest events of ER exit, including the initial exposure of luminal substrates to the cytosol and the spatial organization of this event within the ER, have remained difficult to observe directly. This limitation has hindered a clear separation between initiation of retrotranslocation and downstream processes such as extraction from the ER membrane, ubiquitination, and degradation, leaving fundamental aspects of ER quality control unresolved.

In previous studies, we showed that ERAD substrates accumulate in a specialized, juxtanuclear ER-derived quality control compartment (ERQC) ^19–23^, leading to the proposal that this region functions as a staging ground for ERAD. This interpretation is supported by independent observations of ER reorganization under proteotoxic stress in mammalian cells^24^, as well as the identification of ERAD-related microcompartments in *Chlamydomonas reinhardtii* ^25^. However, these studies relied on indirect or endpoint measurements, and direct experimental evidence linking the initiation of retrotranslocation to the ERQC in mammalian cells has been lacking.

Multiple protein complexes have been proposed as candidates to mediate retrotranslocation. Early models implicated the forward Sec61 translocon as a potential channel for substrate exit^26, 27^, while subsequent studies identified a network of ER luminal and membrane-associated cofactors - including BiP ^28^, calnexin^29, 30^, Sec63p ^28^, protein disulfide isomerase (PDI) ^31^ and the Derlin proteins as contributors to this process ^32, 33^. Particular attention has focused on the Sel1L/HRD1 complex ^34, 35^, which assembles with additional cofactors such as OS-9/XTP3-B, Derlins, and HERP, and has been proposed to function as a central hub for substrate recognition and membrane exit ^4, 14, 36–38^. Alternative models support a tetrameric Derlin-1 channel that recruits p97 for extraction^39–41^ and additional E3 ligases, including gp78/ AMFR, may operate in parallel or substrate-specific pathways ^42^. Membrane context also influences dislocation, with lipid composition and ER factors such as TMUB1 helping to engage p97 ^43^. Moreover, an ER membrane-thinning model has been proposed for HRD1-mediated dislocation, facilitating transfer of substrates to the cytosol, and our recent work implicates oligosaccharyltransferase (OST) as another ER membrane complex with similar activity and acting with HRD1 to promote ERAD ^44^.

Despite these advances, most experimental approaches have so far inferred retrotranslocation from downstream readouts - such as ubiquitination, extraction into the cytosol, or proteasomal degradation - rather than directly monitoring the initiation of cytosolic exposure. As a consequence, it remains unclear where retrotranslocation is initiated within the ER, whether this process is spatially organized, and how early exposure events are mechanistically coupled to subsequent extraction and degradation.

To directly probe the earliest steps of ER exit, we exploited an *Escherichia coli*–derived Biotin Acceptor Peptide (BAP)–Biotin Ligase (BirA) system that enables irreversible biotinylation of luminal protein segments upon their first exposure to the cytosol^45^. Unlike conventional ERAD assays that rely on downstream readouts such as ubiquitination, extraction, or degradation, this approach allows detection of substrates at the moment they become cytosol-accessible, independently of their subsequent fate^46, 47^. Here, we applied this strategy to the well-established ERAD model substrate, H2a^48, 49^. We used BirA-mediated biotinylation as a selective reporter of initial cytosolic exposure of the luminal region of H2a during retrotranslocation initiation. By combining this biochemical readout with high-resolution expansion fluorescence microscopy, we sought to determine whether early retrotranslocation events are spatially organized within the ER, and to directly test whether initiation of ERAD occurs at the ERQC. This approach allows the initiation of retrotranslocation to be interrogated as a spatially and temporally defined event, rather than inferred from downstream extraction or degradation, providing direct evidence linking retrotranslocation to a specific subcellular location.

## Results

### The BAP-BirA system reveals early cytosolic exposure of an ERAD substrate

To monitor early cytosolic exposure during retrotranslocation, we utilized a well-characterized ERAD model substrate, H2a, a type-II membrane protein, isoform of the H2 subunit of the asialoglycoprotein receptor (ASGPR), which is specifically expressed in human hepatocytes ^48^. H2a undergoes cleavage and secretion in hepatocytes; but, when ectopically expressed in other cell types, it remains predominantly uncleaved, retained in the ER, and targeted for ERAD, making it an ideal model substrate for ERAD studies ^49^. Because wild-type H2a can undergo partial proteolytic cleavage, generating a soluble secreted fragment, we used the cleavage-defective H2aG78R mutant^22, 48^ to avoid heterogeneity and ensure that all detected molecules remain competent for ERAD. A Biotin Acceptor Peptide (BAP) tag was added to the luminal C-terminus of H2aG78R, followed by an SV5 epitope tag (H2a-BAP; Fig. 1a). In this configuration, the tagged luminal domain of H2a-BAP becomes accessible to cytosolic BirA upon retrotranslocation, allowing selective detection by BirA-mediated biotinylation. Several non-mutually exclusive mechanisms could, in principle, account for cytosolic exposure of the luminal domain, including passage through a retrotranslocation channel (I), topological inversion via transmembrane domain flipping (II), or complete transfer of the protein into the cytosol (III), potentially facilitated by lipid bilayer thinning (Fig. 1a).

**Figure 1:**
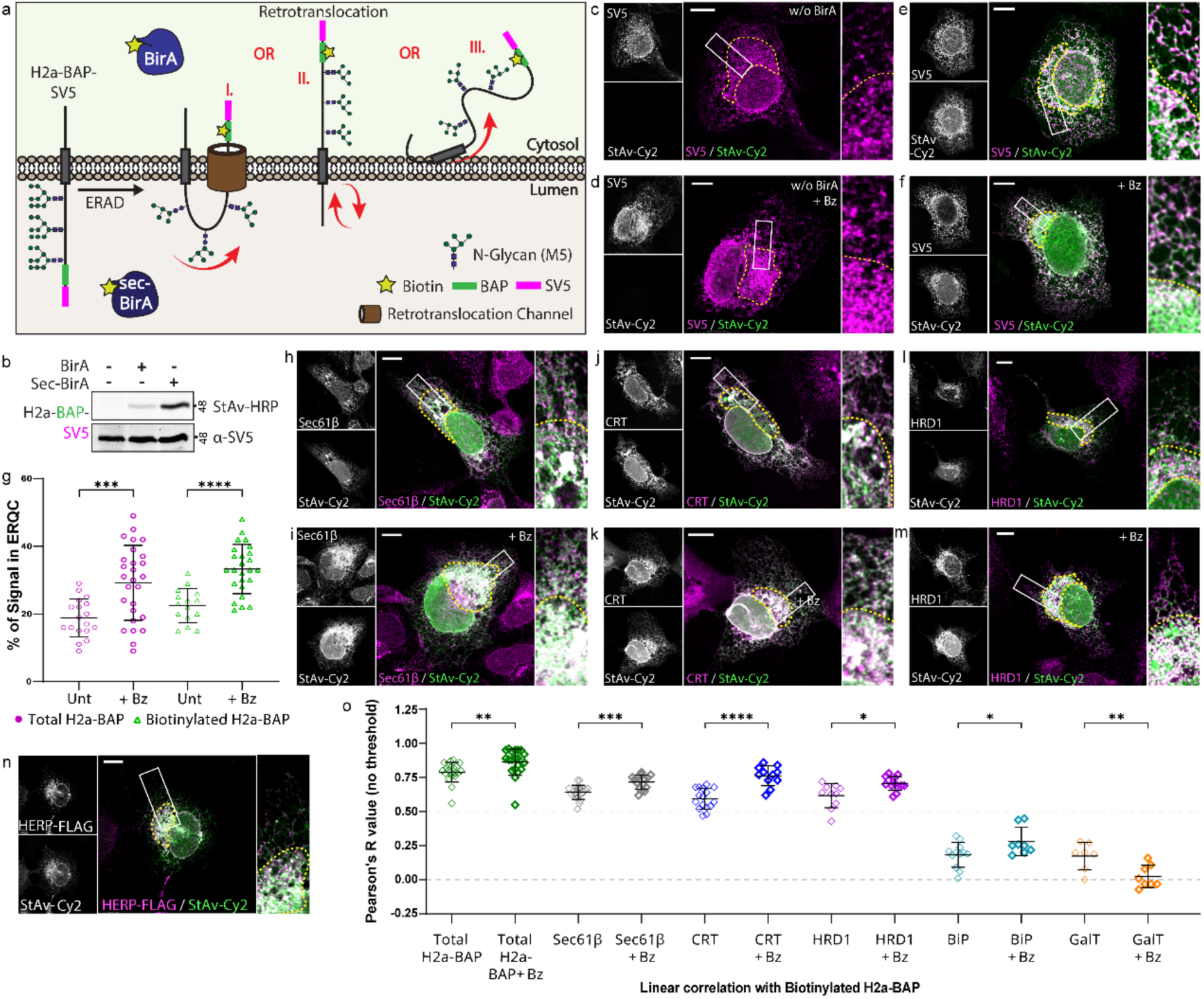
The BAP-BirA system allows visualization of ERAD substrates during initial cytosolic exposure at the ERQC. **a.** Illustration of the ER membrane topology of the model ERAD substrate H2a-BAP-SV5 together with cytosolic BirA and the luminal sec-BirA, highlighting three proposed retrotranslocation mechanisms that allow cytosolic BAP biotinylation. **b.** Western blot of HEK 293 cells expressing H2a-BAP alone or together with BirA or sec-BirA after incubation with Biotin (100µM, 30 min). Biotinylated H2a-BAP was detected with StAv-HRP and total H2a-BAP with α-SV5. **c, d.** Immunofluorescence of U2OS cells expressing H2a-BAP without BirA untreated or upon bortezomib (Bz, 2.5µM, 3 hours) treatment, stained with α-SV5 and StAv-Cy2 to visualize total and background biotin signal; yellow dashed ROIs mark the ERQC and white boxes indicate zoomed regions, shown on the right. **e, f.** Localization of total and biotinylated H2a-BAP in U2OS cells expressing the H2a-BAP – BirA system under basal and Bz-treated conditions. **g.** Quantification of total and biotinylated H2a-BAP in the ERQC relative to whole-cell signal (mean ± SD, n=28; unpaired t-test with Welch’s correction). **h-k.** Colocalization of biotinylated H2a-BAP with endogenous ER/ERQC markers, Sec61β and calreticulin (CRT). **l-n**. Colocalization of biotinylated H2a-BAP with ERAD components, endogenous HRD1 and transfected HERP-FLAG. Scale bars= 10 µm. **o.** Pearson’s R values (Coloc2) for colocalization of biotinylated H2a-BAP with total-H2a-BAP, Sec61β, CRT, HRD1, BiP, and GalT (BiP and GalT from Suppl. Fig. 1) in untreated and Bz-treated cells (unpaired T-test with Welch’s correction).

Consistent with previous work, H2a-BAP displayed the expected topology of a type II membrane protein, confirming correct luminal orientation of the BAP tag ^50^. Western blot analysis of cell lysates (prepared in the presence of N-ethylmaleimide to prevent post-lysis biotinylation) with anti-SV5 to detect total H2a-BAP levels, or with streptavidin linked to HRP (StAv-HRP) to detect only biotinylated molecules, revealed robust biotinylation of H2a-BAP when co-expressed with ER-luminal sec-BirA, whereas only a small fraction of H2a-BAP was biotinylated in the presence of cytosolic BirA (Fig. 1b). This pattern is consistent with only a fraction of H2a-BAP actively undergoing retrotranslocation at any given time, before it is degraded, in contrast to unrestricted access of sec-BirA within the ER lumen. (We later show, through a series of different approaches, that only retrotranslocating H2a-BAP molecules are biotinylated (Fig. 3).)

Immunofluorescence microscopy in U2OS and NIH 3T3 cells showed that H2a-BAP exhibited a reticular ER-like pattern with pronounced enrichment in a distinct juxtanuclear region (Fig. 1c, Suppl. Fig.1a, yellow dotted line). H2a-BAP accumulation in this region was further enhanced upon proteasomal inhibition with bortezomib (Bz) (Fig. 1d), resembling the previously described ER-derived quality control compartment (ERQC) in which ERAD substrates accumulate under conditions of impaired degradation ^19–23^.

We next asked whether retrotranslocating H2a-BAP molecules could be visualized directly using streptavidin-conjugated fluorophores to detect BirA-mediated biotinylation in the cytosol. This would reveal actively-retrotranslocating molecules, before their proteasomal degradation. Indeed, this approach produced no detectable non-specific labeling (Fig. 1c-d) and revealed a striking enrichment of biotinylated H2a-BAP within the ERQC upon proteasome inhibition, as well as substantial localization to this compartment under basal conditions (Fig. 1e-f; Suppl. Fig. 1b). Biotinylated H2a-BAP largely colocalized with total H2a-BAP within the ERQC, whereas in peripheral ER regions the streptavidin signal was comparatively reduced relative to total H2a-BAP. Quantitative analysis confirmed a significant enrichment of retrotranslocating molecules at the ERQC following proteasomal inhibition (Fig. 1g).

Consistent with previous observations, the ERQC was enriched in ERAD-related components. Sec61β and calreticulin (CRT), previously reported to relocalize to the ERQC upon ERAD substrate accumulation ^19, 20^, showed pronounced colocalization with biotinylated H2a-BAP within this compartment (Fig. 1h-k, o, Suppl. Fig. 1e-f). In contrast, the abundant ER chaperone BiP was largely excluded from the ERQC, in agreement with previous reports^19, 20^, and did not show practically any colocalization with retrotranslocating H2a-BAP (Fig. 1o; Suppl. Fig. 1c). Although the ERQC is located in the centrosomal region of the cell ^19, 20, 51^, biotinylated H2a-BAP did not colocalize with the Golgi marker GalT-YFP, indicating that these compartments remain distinct (Fig. 1o; Suppl. Fig. 1d). The ERAD lectin OS-9^52^, an established ERQC component associated with the HRD1 complex^53^, also showed strong colocalization with retrotranslocating H2a-BAP (Suppl. Fig. 1h), further supporting the existence of a staging ground for retrotranslocation and ERAD at the ERQC.

We next examined the localization of retrotranslocating H2a-BAP relative to endogenous HRD1, a central ERAD-associated ubiquitin ligase^38, 52–55^. Biotinylated H2a-BAP showed strong colocalization with HRD1 at the ERQC (Fig. 1l, m, o). HERP, which has been implicated in recruitment and stabilization of ERAD machinery at the ERQC^22^, similarly colocalized with retrotranslocating H2a-BAP, predominantly at this compartment (Fig. 1n; Suppl. Fig. 1g).

Altogether, these findings identify the ERQC as a major site where ERAD substrates engage the ERAD machinery and undergo initial cytosolic exposure during retrotranslocation. They also establish the BAP-BirA system as a valuable tool for the visualization of ERAD substrates at the onset of retrotranslocation.

### Higher resolution imaging of the subcellular sites of retrotranslocation

We next exploited the H2a-BAP – BirA system to localize retrotranslocation sites at higher spatial resolution using protein-retention expansion microscopy (see Methods). In untreated U2OS cells, total H2a-BAP (anti-SV5) displayed a reticular ER distribution, with a denser juxtanuclear domain consistent with the ERQC (Fig. 2a-f; ERQC outlined in Fig. 2c; Suppl. Fig.2a, b). Whereas standard confocal microscopy (Fig. 1, Suppl. Fig. 1) portrays this region as a compact blurred structure, expansion microscopy physically separated closely apposed membranes and increased effective lateral and axial resolution, revealing discrete, stacked ER meshes within the ERQC. These structures were readily apparent in sum-intensity z-projections (Fig. 2c-d) and in temporal color-coded (Fire LUT) hyperstacks representing successive optical cellular sections across the z-axis (Fig. 2e-f). Co-imaging total and biotinylated H2a-BAP in untreated cells revealed broad ER distribution of the total protein, with selective enrichment of the retrotranslocating species in the ERQC relative to the peripheral ER (Fig. 2g-l; Suppl. Fig. 2c, d), reinforcing and extending our previous observation that retrotranslocation occurs predominantly at the ERQC. Proteasome inhibition (with lactacystin, Lac) further intensified this phenotype: total H2a-BAP was still seen throughout the ER, but accumulated strongly in the ERQC, which appeared thicker and more densely layered by expansion (Fig. 2m-r; Supp. Fig. 2e, f). Under these conditions, retrotranslocating H2a-BAP molecules became even more concentrated at the ERQC and traced a multilayered, mesh-like network resolvable only after expansion (Fig. 2s-x; Suppl. Fig. 2g, h), a structural organization not discernible by conventional confocal microscopy. These findings provide direct visualization of a dense, multilayered mesh-like ER architecture within the mammalian ERQC and directly associate this architecture with sites of active retrotranslocation (Fig. 2x, y).

**Figure 2:**
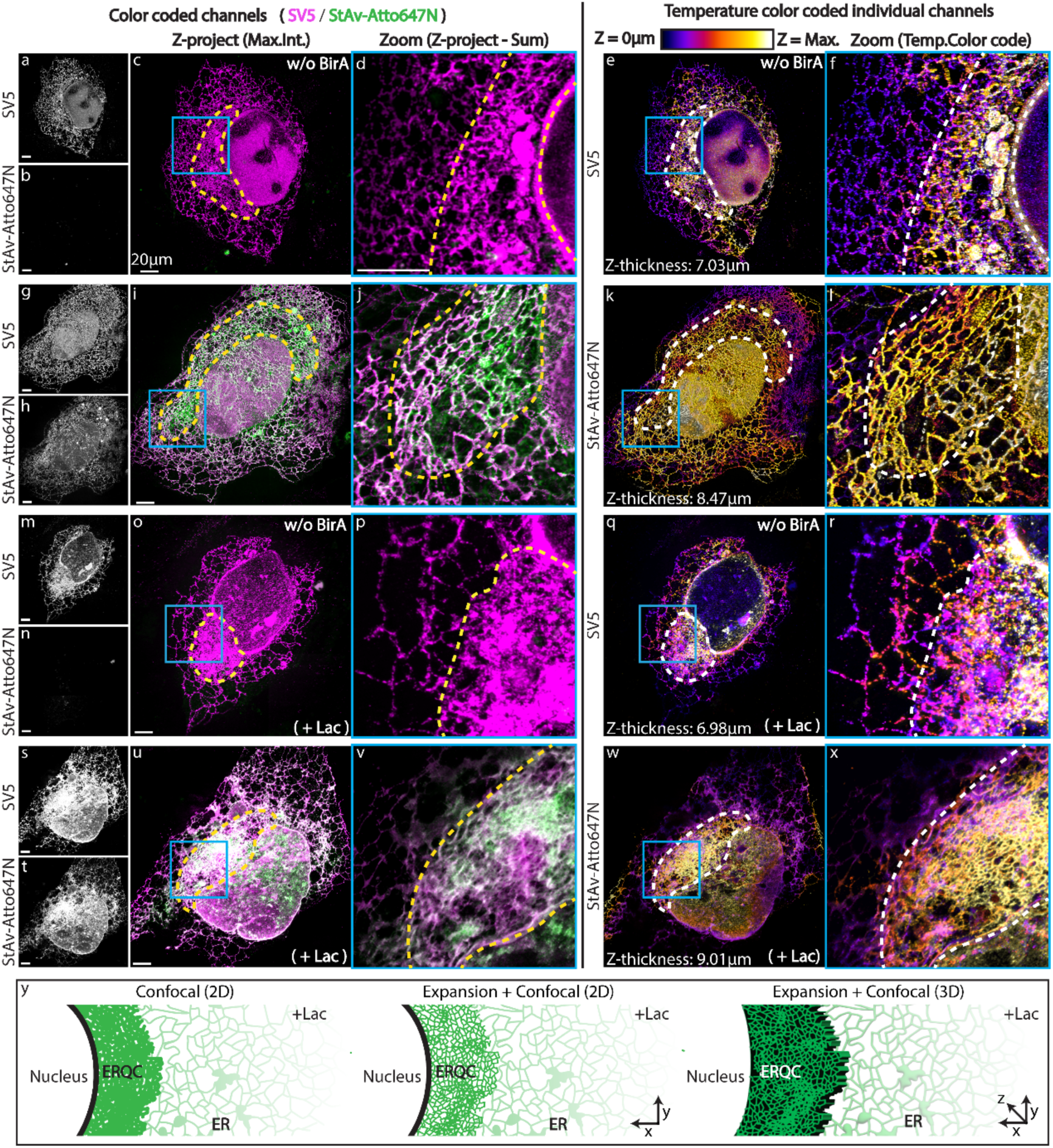
Expansion microscopy reveals a multilayered ERQC architecture enriched in retrotranslocating substrates. U2OS cells expressing H2a-BAP alone or together with BirA were left untreated or treated with the proteasome inhibitor Lactacystin (Lac, 25 µM, 3 hour) and processed for expansion microscopy. After gel embedding and expansion, gels were incubated with α-SV5 or StAv-Atto647N to detect total and biotinylated H2a-BAP, cut to imaging-sized pieces, mounted between 25-mm coverslips and imaged on a Leica SP8 confocal microscope using a 63× oil objective in Lightning mode with water as mounting medium. Z-stacks were processed in ImageJ to generate maximum-intensity (**c, i, o, u**) or sum-intensity (**d, j, p, v**) projections, and the indicated individual channels were displayed as temporally color-coded stacks using a Fire LUT (**e, k, q, w**) with corresponding zooms (**f, l, r, x**). Yellow/white dashed lines indicate ERQC regions of interest and blue boxes mark zoomed areas. Scale bars= 20 µm. Images are representative of three independent experiments. **y.** Scheme comparing ERQC resolution in conventional confocal images (left) and in expanded samples imaged in 2D (middle) or 3D (right), illustrating the 3D volume of the ERQC relative to the 2D ER network.

Building on prior work implicating COPII-dependent delivery of ERAD substrates to the ERQC and COPI-mediated recycling back to the peripheral ER^56^, we wondered whether perturbation of these pathways would affect H2a-BAP retrotranslocation. Inhibition of COPII transport with dominant-negative Sar1(T39N) reduced the ratio of biotinylated H2a-BAP to total H2a-BAP to about half, whereas conversely, inhibiting COPI with dominant-negative ARF1(T31N) increased this ratio approximately 2-fold (Suppl. Fig. 3a, b). These effects are consistent with the ERQC functioning as a principal site for H2a-BAP retrotranslocation and degradation (Suppl. Fig. 3c).

### The BAP–BirA system captures the onset of H2a retrotranslocation

To validate that BirA-dependent biotinylation reports cytosolic exposure of H2a-BAP molecules, we first compared its biotinylation by the cytosolic BirA and the ER-luminal sec-BirA to a panel of BAP-tagged proteins with distinct topologies (Fig. 3a). For H2a-BAP, cytosolic BirA labeled 24.6% of the population relative to sec-BirA (Figs. 1b, 3c), consistent with detection of a subfraction which is transiently exposed to the cytosol during retrotranslocation. In contrast, BAP-H2a, in which the BAP tag is N-terminal and constitutively cytosolic, showed robust biotinylation by BirA, and a comparable signal with sec-BirA (Figs. 3b, c), likely reflecting incomplete ER targeting of sec-BirA, resulting in a residual cytosolic pool. A soluble cytosolic control, BAP-p97, exhibited a labeling pattern similar to that of BAP-H2a, confirming efficient biotinylation of cytosolic BAP-tagged proteins by BirA and partial cytosolic access of sec-BirA (Figs. 3b, c). As a luminal control, BiP-BAP showed limited BirA labeling (15.3%) (Figs. 3b, c), in line with reports that a minor fraction of BiP can appear in the cytosol under specific conditions ^57, 58^, potentially enhanced by overexpression. Together, these controls with varied topology validate H2a-BAP – BirA as a selective reporter of cytosolic exposure.

**Figure 3:**
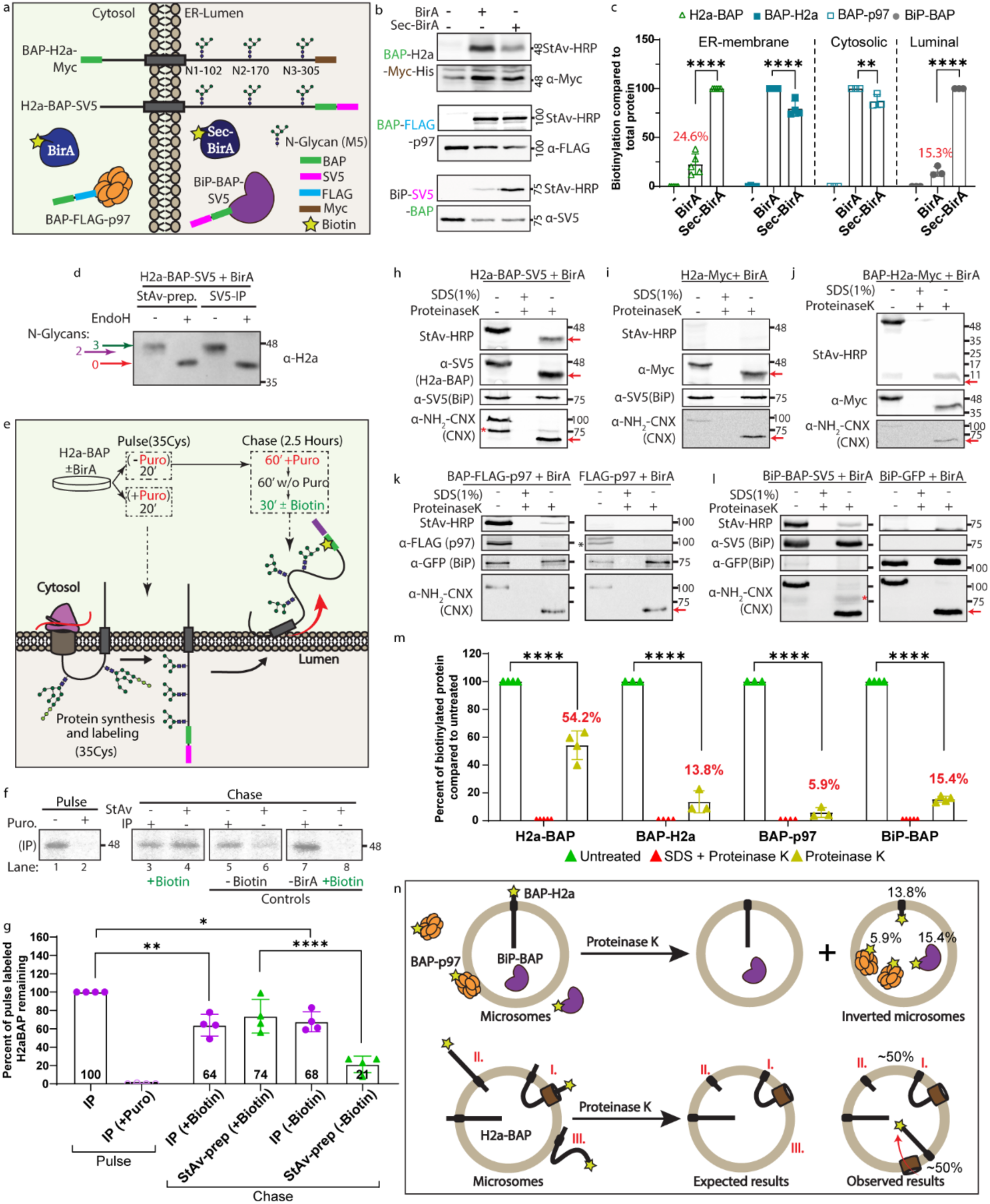
BirA-mediated labeling detects post-translational cytosolic exposure of H2a-BAP during retrotranslocation. **a.** Illustration showing the localization and membrane topology of the BAP-tagged constructs. **b.** Western blot analysis of HEK 293 cells expressing BAP-tagged constructs alone or with BirA or sec-BirA after incubation with biotin (100 µM, 30 min); biotinylated BAP-H2a-Myc-His, BAP-FLAG-p97 and BiP-SV5-BAP were detected with StAv-HRP and total protein with tag-specific antibodies. **c.** Quantification of signals from Figs. 1b and 3b, normalized to sec-BirA biotinylation for H2a-BAP and BiP or to BirA biotinylation for cytosolic controls (BAP-H2a and p97), shown as mean of ≥3 experiments ± SD **d.** Immunoprecipitated total H2a-BAP and StAv-precipitated biotinylated H2a-BAP, ± Endo H, were compared by Western blot with α-H2a antibody (against N-terminus); colored arrows indicate the number of N-glycans, according to the differences in migration of H2a glycosylation site mutants (Suppl. Fig. 4a). **e.** Schematic of the pulse-chase experiment (20-min [³⁵S]-cysteine pulse, 2.5-hour chase). **f.** Total H2a-BAP (anti-H2a IP) and biotinylated H2a-BAP (StAv pull-down) from the pulse-chase labeling were analyzed by SDS-PAGE; puromycin was present during the first 1 hour of chase, biotin was added at the end of the chase (100 µM, 30 min) and controls lacked biotin or BirA. **g.** Quantification of pulse-chase signals (mean of 4 experiments ± SD). **h-l.** Protease protection assays in HEK 293 cells co-expressing BAP-tagged constructs (H2a-BAP-SV5, BAP-H2a-Myc, BAP-p97-FLAG, and BiP-BAP-SV5) with BirA after biotin incubation (100 µM, 30 min.); homogenates were left untreated, treated with proteinase K, or with proteinase K after SDS lysis, and biotinylated proteins were detected with StAv-HRP while total proteins (indicated in parentheses) were detected with tag-specific antibodies. BiP-BAP-SV5 or BiP-GFP served as luminal protection controls and N-terminal CNX as a cytosolic sensitivity control; red arrows mark cleavage of cytosolic domains and the red asterisk in CNX panels marks a cross-reactive BiP-BAP-SV5 signal. **m.** Protease protection signals were quantified, normalized to untreated samples and plotted as mean of ≥3 experiments ± SD. Statistics were performed using ordinary two-way ANOVA with Tukey’s multiple-comparisons test. **n.** Illustration summarizing occasional microsome inversions and proteinase K sensitivity of H2a-BAP in three retrotranslocation models (I-III).

Because the BAP tag is located at the C-terminus of H2a, the probability of its exposure to cytosolic BirA during co-translational forward-translocation of H2a-BAP is expected to be rare. To test whether BirA labeling occurs exclusively after ER entry and retrotranslocation, rather than during forward-translocation, we examined N-glycan occupancy of biotinylated H2a-BAP (Fig. 3d). Biotinylated H2a-BAP (streptavidin pull-down) and total H2a-BAP (SV5 - IP) were resolved by SDS–PAGE, with or without treatment with Endo H (which removes all N-linked high mannose sugar chains). The biotinylated molecules displayed predominantly full N-glycan occupancy, indicating that these molecules had completed forward translocation into the ER lumen before cytosolic exposure (Fig. 3d). Minor underglycosylated species, corresponding to incomplete modification of a single glycosylation site, were detected at similarly low levels in both the biotinylated and total H2a-BAP pools. Analysis of glycosylation-site mutants confirmed that site 305 is the most frequently under-modified position in H2a, accounting for the residual underglycosylated species observed. The H2a Δ120/170 mutant (retaining only site 305) showed ∼40% non-glycosylated protein, whereas the Δ305 mutant lacked any underglycosylated band (Suppl. Fig. 4a). Together, these data indicate that BirA-dependent labeling of H2a-BAP occurs essentially after ER entry, luminal N-glycosylation, and initiation of retrotranslocation, rather than during co-translational forward translocation.

To further define the timing of the biotinylation event, we performed a delayed-biotin pulse-chase analysis using metabolic labeling with [^35^S]-cysteine (Fig. 3e). Following the radioactive pulse, cells were chased for 2.5 hours. During the first hour of the chase, puromycin was added to block any residual [^35^S]-labeled protein synthesis. After a recovery period of 1 hour without puromycin, exogenous biotin was added late in the chase for 30-minutes. Biotinylated [^35^S]-labeled H2a-BAP (detected by streptavidin pull-down, StAv) and total [^35^S]-labeled H2a-BAP (immunoprecipitated, IP) were detected and analyzed. Under these conditions, robust biotinylation of pre-existing [^35^S]-labeled H2a-BAP was observed (Fig. 3f, lane 4), comparable to total [^35^S]-H2a-BAP (lane 3), whereas control conditions lacking BirA or exogenous biotin showed little or no detectable biotinylation signal (lanes 6,8, quantification in Fig.3g). Another control included puromycin during the pulse, which showed its prevention of any [^35^S] labeling (lane 2). Since no newly synthesized H2a-BAP molecules were [^35^S]-labeled during the biotin incubation, these results demonstrate that biotinylation of pre-existing [^35^S]-labeled H2a-BAP molecules by BirA occurs post-translationally, after ER entry, and marks H2a-BAP molecules undergoing retrotranslocation rather than forward translocation.

We next asked whether biotinylated H2a-BAP molecules remain exposed to the cytosol before their degradation. Proteinase K protease-protection assays performed on microsomes revealed that, surprisingly, approximately half of the biotinylated H2a-BAP population was protected from proteolysis, yielding a protected fragment (corresponding to the luminal and transmembrane domains) (54.2% protected; Fig. 3h, m). A comprehensive set of control substrates confirmed assay specificity and enabled estimation of microsome inversion, which accounted for only a minor fraction of the protected signal (Fig. 3i-m). Non-BAP-tagged H2a-Myc coexpressed with BirA produced no biotinylation signal but underwent complete proteolytic cleavage, as expected (Fig. 3i); BAP-H2a (with the N-terminal BAP tag constitutively cytosolic) lost nearly all biotin signal upon protease treatment, as expected, yet showed minor protection of a ∼10 kD N-terminal cytosolic fragment (13.8% protected; Fig. 3j, m); biotinylated cytosolic BAP-p97 showed minimal protection (5.9%; Fig. 3k, m); whereas the biotinylated luminal control BiP-BAP exhibited 15.4% protection (Fig. 3l, m). As we had seen that 15.3% of total Bip-BAP was biotinylated (Fig. 3c), and 15.4% of biotinylated BiP-BAP was protease-protected (Fig. 3m), this implies that only ∼2.35% of total BiP is both biotinylated and protected. This value is consistent with inversion of a minor fraction of microsomes during lysis (Fig 3n). As microsome inversion therefore accounts for only a minority of protected molecules, the substantially larger protected fraction observed for biotinylated H2a-BAP is consistent with regression or back-sliding of the C-terminus back to the ER-lumen after transient cytosolic exposure and biotinylation. Given that the cytosolic population is readily ubiquitinated and degraded by proteasomes in the cell, while the luminal population is protected, this latter population is expected to be over-represented at steady state. Consistently, biotinylated H2a-BAP showed negligible polyubiquitination under basal conditions, which increased substantially upon proteasome inhibition (anti-Ubiquitin; Fig. 6a). Together, these results suggest that H2a retrotranslocation involves transient cytosolic exposure followed by partial regression prior to ubiquitin-dependent commitment to degradation. This backsliding behavior favors a mechanism via a channel-mediated retrotranslocation, (Fig. 1a - Model I), because the reversal of a topology inversion (Model II) or of a complete cytosolic transfer (Model III) would be very energy-demanding. In this mechanism, incomplete extraction permits back-sliding before stabilization by polyubiquitination, consistent with ratcheting models previously proposed for ERAD substrates^36, 59^.

### Retrotranslocation primes extraction-dependent deglycosylation and degradation of H2a-BAP

To better define where retrotranslocating H2a-BAP resides and how it is processed, we fractionated HEK 293 derived microsomes on an iodixanol gradient optimized to resolve ER subdomains ^22, 56, 60^ (Fig. 4a-c). Organelle markers behaved as expected, Cab45 in light Golgi-derived fractions (1-2), RPL26 (a ribosomal subunit) in heavy rough-ER fractions (8-11), and ER membrane proteins (CNX, HRD1, Sec61β) distributed across intermediate fractions (3-8), confirming appropriate gradient resolution (Fig. 4d). Under basal conditions, retrotranslocating H2a-BAP (detected by StAv-HRP) co-migrated with total H2a-BAP (detected by anti-SV5) across ER fractions, indicating that retrotranslocating species remain membrane-associated prior to degradation (Fig. 4a, d). Proteasome inhibition with bortezomib (Bz) caused a modest shift of both total and retrotranslocating H2a-BAP toward heavier fractions (7-9) (Fig. 4b, d), consistent with accumulation in a denser ER quality-control compartment (ERQC), as previously observed for WT H2a and other ERAD substrates^22^. In addition, a faster-migrating H2a-BAP species appeared in light density membrane and soluble fractions (1-3) upon proteasome inhibition (Fig. 4b, red arrows), but was scarcely detectable under basal conditions (Fig. 4a, d). This species comigrated with deglycosylated H2a-BAP generated by Endo H or PNGase F treatment (Suppl. Fig. 4b, c). These results are in line with H2a-BAP deglycosylation followed by fast degradation, whereas the deglycosylated species accumulates in the cytosol when proteasomes are inhibited.

**Figure 4:**
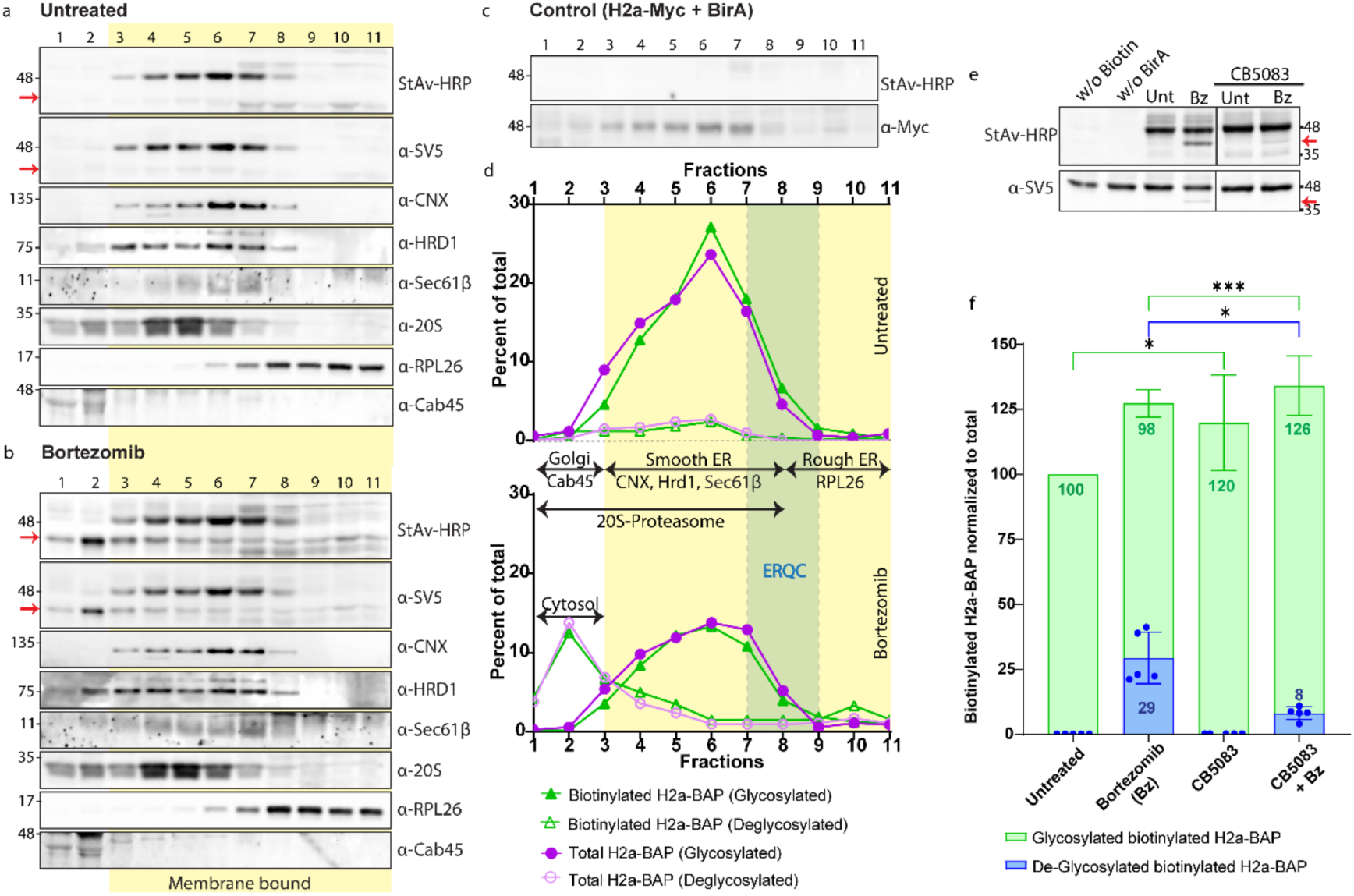
Retrotranslocating H2a-BAP remains membrane-associated prior to p97-dependent extraction, deglycosylation and degradation. **a, b.** H2a-BAP – BirA expressing HEK 293 cells, left untreated (a) or treated with Bz (2.5 µM, 3 hours) (b), were homogenized and post-nuclear supernatants were layered on iodixanol step gradients (10, 16, 22, 28 and 34%) and centrifuged at 24,000 r.p.m. for 16 hours. 1 mL fractions were collected from the top and analyzed by Western blot with StAv-HRP to detect biotinylated H2a-BAP and with α-SV5 (or α-Myc) for total H2a-BAP, together with organelle markers α-CNX, α-HRD1, α-Sec61β, α-20S, α-RPL26 and α-Cab45; the deglycosylated H2a-BAP species is indicated by a red arrow. **c**. H2a-Myc lacking the BAP tag was co-expressed with BirA as a negative control for biotinylation. **d.** For each protein, the signal in every fraction was expressed as a percentage of total and plotted versus fraction number; one representative experiment of three is shown. Yellow shading marks fractions enriched in membrane-bound organelles, with fractions 1–2 containing soluble proteins and low-density organelles (Golgi), and green shading marks the ERQC peak. **e.** HEK 293 cells expressing H2a-BAP - BirA were treated with the p97 inhibitor CB5083 (2.5 µM, 3 hours), alone or with Bz (2.5µM, 3 hours), and analyzed by Western blot. **f.** Glycosylated and deglycosylated biotinylated H2a-BAP species (from (e)) were quantified, normalized to total H2a-BAP (α-SV5) and expressed relative to untreated cells (mean of 5 experiments ± SD; ordinary two-way ANOVA with Tukey’s multiple-comparisons test).

To test whether extraction from the membrane is required for deglycosylation, we inhibited (using CB5083) the AAA ATPase p97, known to extract membrane proteins during ERAD (Fig. 4e). Interestingly, p97 inhibition did not decrease the abundance of retrotranslocating H2a-BAP molecules detected by BirA-mediated biotinylation (StAv-HRP); instead, a modest increase was observed. Total H2a-BAP levels were also moderately increased by p97 inhibition, consistent with impaired degradation. However, no deglycosylated H2a-BAP was observed after p97 inhibition. Co-inhibition of p97 and the proteasome caused an approximately 73% reduction of the deglycosylated species compared to proteasome inhibition alone (Fig. 4f). These results indicate that p97-mediated extraction is required for deglycosylation and subsequent degradation, whereas initial cytosolic exposure during retrotranslocation does not depend on p97 activity.

We next asked whether degradation proceeds after complete release of the ERAD substrate into the cytosol as a soluble species or while it remains membrane-associated. Using carbonate extraction, followed by iodixanol gradient fractionation, we distinguished integral membrane proteins from species peripherally associated to the membrane. As controls, ribosomes (RPL26) and eIF2α (cytosolic proteins that partially bind membranes) shifted to light (soluble) fractions upon SDS lysis or carbonate treatment (Fig. 5b), whereas integral ER proteins (CNX, HRD1) remained membrane-associated after carbonate extraction but were solubilized by SDS (Fig. 5a, b). Under basal or proteasome-inhibited conditions, both glycosylated retrotranslocating (StAv-HRP) and total (SV5) H2a-BAP behaved as integral membrane proteins (Fig. 5a, c, e). In contrast, deglycosylated H2a-BAP detected under basal conditions was membrane-associated but not integrated, becoming soluble upon carbonate extraction (Fig. 5a, d, f). Upon proteasome inhibition, deglycosylated H2a-BAP was predominantly soluble with a small membrane-associated fraction remaining (Fig. 5a, f). This suggests that deglycosylation starts on membrane-associated (but not integrated) species that are then sent to the cytosol for proteasomal degradation. Proteasomes were distributed mainly across medium density membrane fractions and shifted to soluble fractions upon SDS lysis, as expected, whereas carbonate extraction also shifted their peak toward soluble fractions but left a substantial membrane-tethered population (Figs. 4a, b, 5b). Under proteasome inhibition, partial co-sedimentation of proteasomes with deglycosylated H2a-BAP in mid-density fractions possibly reflects substrate-proteasome assemblies at the membrane.

**Figure 5:**
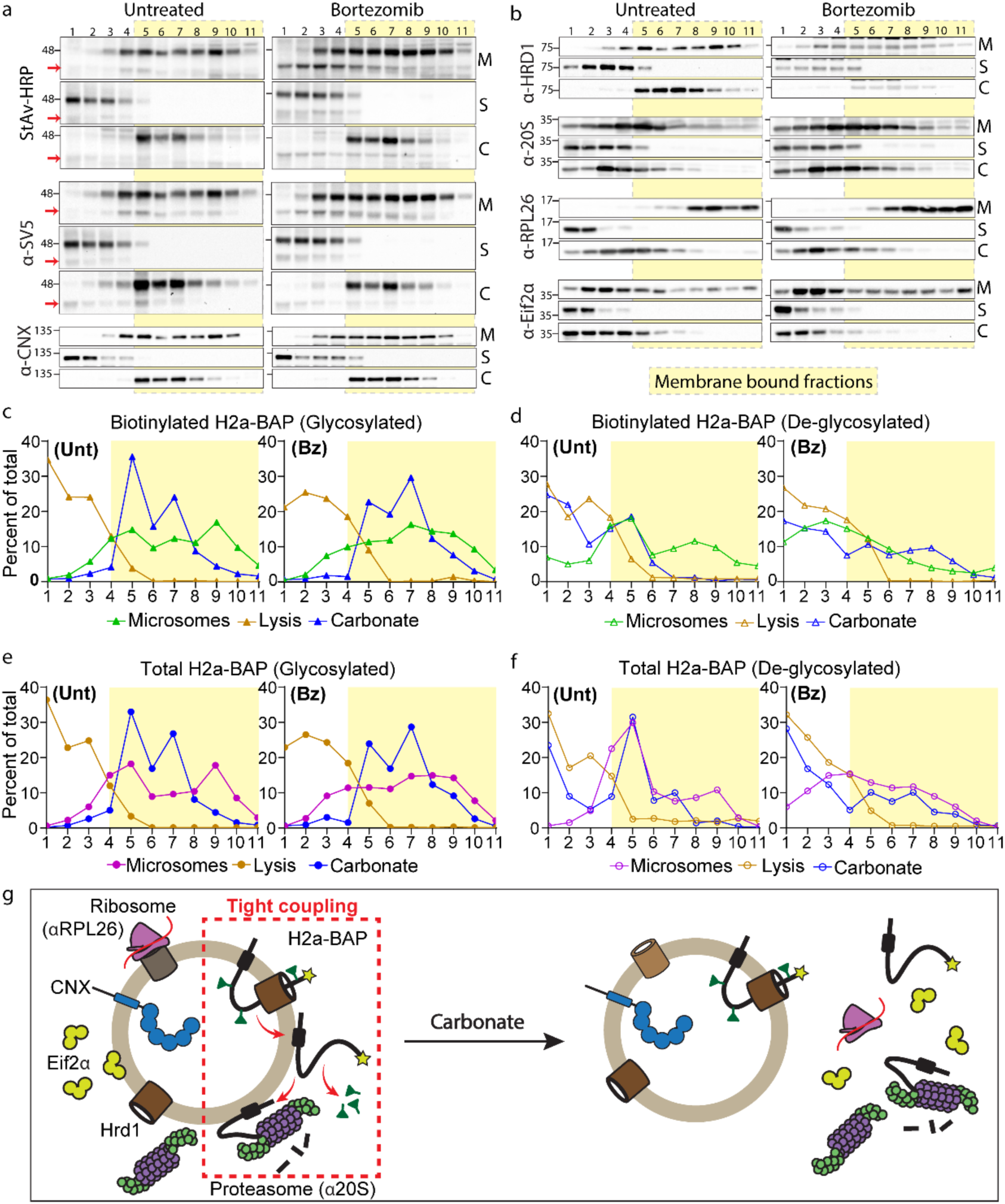
Deglycosylation and proteasomal engagement occur at the membrane-cytosol interface during ERAD. H2a-BAP – BirA expressing HEK 293 cells were processed as in Fig. 4a but using iodixanol gradients optimized to achieve higher resolution of low and medium density fractions (10%, 13%, 16%, 20% and 24%) under three microsome conditions: untreated microsomes (M), SDS/Triton X-100/NaDOC lysis (S) or Na₂CO₃ extraction (C). **a.** 1-mL fractions were collected and immunoblotted for biotinylated H2a-BAP (StAv-HRP), deglycosylated species are indicated by red arrows), total H2a-BAP (α-SV5) and calnexin (CNX). **b.** Parallel blots were probed for endogenous HRD1 and cytosolic or cytosol-facing membrane proteins (α-20S, α-RPL26 and α-eIF2α). **c-f.** Signals for biotinylated, total, glycosylated and deglycosylated H2a-BAP were quantified for each condition (M, S, C), expressed as percentage of total signal and compared between untreated and Bz-treated cells; one representative experiment of three is shown. **g.** Scheme of alkaline carbonate extraction release of peripherally membrane-associated proteins from microsomes while retaining integral ER membrane proteins.

Together, these observations support a tightly coupled, membrane-proximal pathway in which retrotranslocation, ubiquitination, p97-dependent extraction, cytosolic deglycosylation, and proteasomal degradation proceed in close spatial and temporal succession at the ER membrane/ERQC interface (Fig. 5g). p97 is required to extract the substrate and enable deglycosylation, and under basal conditions no stable soluble retrotranslocated intermediate accumulates in the cytosol, whereas it does upon proteasome inhibition, implying immediate degradation upon detachment from the membrane.

### The H2a-BAP – BirA system resolves how pathway interference alters retrotranslocation and downstream processing

We leveraged the H2a-BAP – BirA system, in which biotinylation reports cytosolic exposure of retrotranslocating H2a-BAP molecules, to dissect how perturbations in ER quality control influence retrotranslocation and downstream ubiquitination.

To examine the role of early N-glycan processing, we acutely inhibited α-glucosidase I/II using NB-DNJ. As expected, H2a-BAP exhibited a slight electrophoretic mobility shift, consistent with blocked glucose trimming (Fig. 6a). This perturbation caused only a modest change in the fraction of biotinylated H2a-BAP relative to total H2a-BAP (Fig. 6a, b). Streptavidin pulldown followed by anti-ubiquitin immunoblotting revealed no detectable change in polyubiquitination of the retrotranslocating pool (Fig. 6a, c). Proteasome inhibition together with NB-DNJ treatment increased substrate accumulation, but did not further alter the relative extent of retrotranslocation or polyubiquitination beyond that observed with Bz alone (Fig. 6a-c). These results indicate that glucose trimming is dispensable for retrotranslocation and downstream ubiquitination of H2a-BAP, consistent with glycan-independent ERAD routes operating in parallel with canonical lectin-dependent pathways ^61, 62^.

**Figure 6.**
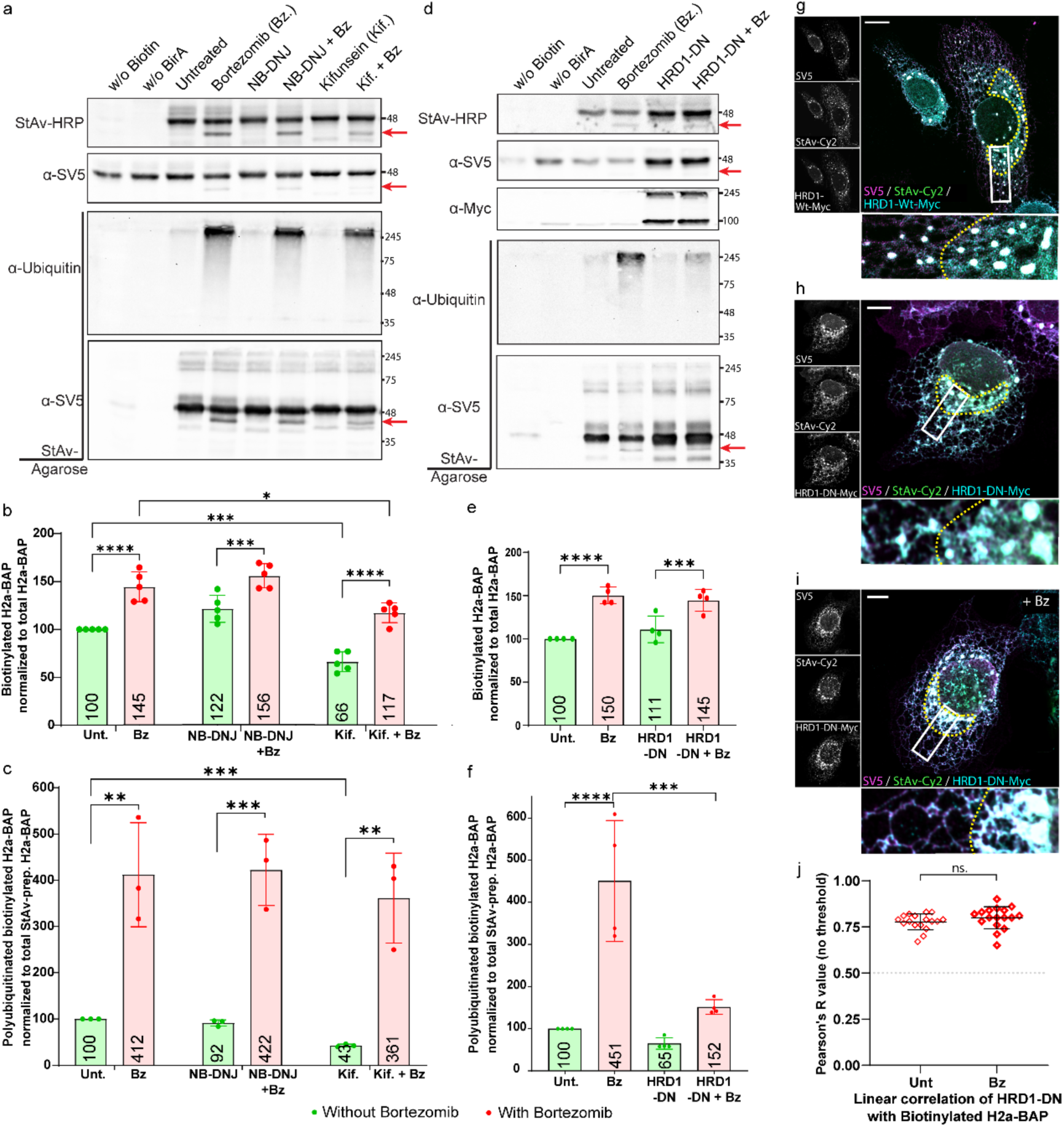
Mannose trimming promotes entry into the retrotranslocation pathway, whereas the E3 ligase HRD1 acts downstream of cytosolic exposure. **a.** HEK 293 cells expressing H2a-BAP - BirA were left untreated or treated for 3 hours with Bz (2.5µM), NB-DNJ (200µM) or Kifunensine (Kif, 100µM), alone or with Bz, and lysates were analyzed for total (α-SV5) and biotinylated H2a-BAP (StAv-HRP) or subjected to StAv-agarose pulldown to detect precipitated (α-SV5) and ubiquitylated H2a-BAP (α-Ubiquitin). Deglycosylated species are indicated by red arrows. **b, c.** Biotinylated H2a-BAP and polyubiquitylated biotinylated H2a-BAP were quantified and normalized to total or StAv-precipitated H2a-BAP, respectively. **d.** HEK 293 cells co-expressing H2a-BAP - BirA with dominant-negative HRD1 (HRD1-DN) were analyzed under basal or Bz-treated conditions using the same biochemical readouts as in (a). **e, f.** Quantification of biotinylated and polyubiquitylated H2a-BAP (from d). Statistics were performed using ordinary two-way ANOVA with Tukey’s multiple-comparisons test. **g-i.** U2OS cells co-expressing H2a-BAP - BirA with HRD1-WT or HRD1-DN, with or without Bz, were stained for total and biotinylated H2a-BAP and HRD1/HRD1-DN (α-Myc) to visualize the impact of HRD1 activity and proteasome inhibition on H2a-BAP localization. **j.** Pearson’s R values (Coloc2) for colocalization of biotinylated H2a-BAP with HRD1-DN under untreated and Bz-treated cells (unpaired T-test with Welch’s correction). Scale bars= 10 µm.

By contrast, inhibition of ER class-I α-mannosidases with kifunensine (Kif) ^63^ strongly impaired degradation of H2a-BAP and markedly affected retrotranslocation, reducing substantially the biotinylated H2a-BAP fraction (Fig. 6a, b). Polyubiquitination of the retrotranslocating pool was also substantially reduced (Fig. 6a, c), placing mannose trimming upstream of productive engagement with the retrotranslocation and ubiquitination machineries. Co-inhibition with Kif and Bz partially restored the levels of polyubiquitination of the residual retrotranslocated molecules while retrotranslocation itself remained reduced. Together, these observations suggest that mannose trimming promotes efficient entry into the retrotranslocation pathway and subsequent ubiquitination, rather than acting directly on ubiquitin chain assembly.

We next examined the role of the E3 ligase HRD1, previously implicated in ERAD of WT H2a. Expression of a dominant-negative RING-inactivated HRD1 mutant (HRD1-DN), caused robust accumulation of total H2a-BAP (Fig. 6d), but did not measurably reduce retrotranslocation efficiency compared to basal conditions or under proteasome inhibition, as assessed by BirA-mediated biotinylation (Fig. 6d, e). As expected, polyubiquitination of the retrotranslocating pool was significantly reduced but not abolished, consistent with residual activity of endogenous HRD1 or compensatory E3 ligases (Fig. 6d, f). Imaging corroborated these results: overexpressed HRD1-WT formed colocalized puncta with H2a-BAP (Fig. 6g; Suppl. Fig. 5a), likely reflecting oligomerization upon overexpression. In contrast, HRD1-DN assembled punctate aggregates along ER tubules, distorted local ER architecture, and concentrated in the ERQC where it strongly colocalized with retrotranslocating H2a-BAP; this sequestration was further accentuated by proteasome inhibition (Fig. 6h-j; Suppl. Fig. 5b). Altogether, HRD1 activity appears dispensable for H2a-BAP retrotranslocation but is required for efficient post-cytosolic exposure polyubiquitination and degradation; defective HRD1 nucleates aberrant assemblies that trap substrates in the ERQC.

Finally, we assessed the contribution of the cytosolic SCF-type Cullin-RING E3 ligase SCF^FBS2^, previously implicated in ERAD of WT H2a^64^, and known to recognize high-mannose glycans. WT FBS2, the F box subunit of this ligase, showed diffuse cytosolic localization with modest enrichment at the ERQC and minor if any colocalization with retrotranslocating H2a-BAP, which increased upon proteasome inhibition (Suppl. Fig. 5c, d), consistent with convergence of luminal and cytosolic ERAD factors in this compartment. Colocalization analysis supported this redistribution: thresholded Pearson’s R (above autothreshold) was negative under basal conditions but became positive upon proteasome inhibition (Suppl. Fig. 5e). In contrast, a dominant-negative FBS2 variant (ΔFBS2, lacking the F-box domain, but glycan-binding competent) formed perinuclear puncta that strongly colocalized with retrotranslocating H2a-BAP while depleting the normal cytosolic FBS2 pool (Suppl. Fig. 5f). These results support a model in which SCF^FBS2^ acts downstream of retrotranslocation to facilitate ubiquitin chain assembly, rather than facilitating the retrotranslocation step itself.

Altogether, these results resolve discrete control points in ERAD of H2a-BAP: *(i)* mannose trimming promotes efficient engagement of the retrotranslocation machinery; *(ii)* HRD1 and SCF^FBS2^ act predominantly downstream of the initial cytosolic exposure to drive ubiquitination; and *(iii)* the ERQC functions as a spatial hub that coordinates glycan-sensing, retrotranslocation, ubiquitination, membrane-extraction and proteasomal degradation.

## Discussion

Retrotranslocation is a central step in ER proteostasis. Its failure leads to misfolded-protein accumulation and ER stress ^5–7^, and it is exploited by bacterial toxins to gain access to the cytosol ^8–14^. Despite its importance, the site and timing of retrotranslocation initiation in mammalian cells have remained unclear. In this study, we provide direct experimental evidence that retrotranslocation initiation during ERAD is a spatially and mechanistically defined process that occurs at a specialized ER-derived quality control compartment (ERQC). By combining a biotin-based reporter of transient cytosolic exposure with high-resolution expansion microscopy, we visualize ERAD substrates at the moment they first engage the retrotranslocation machinery, independently of subsequent extraction, ubiquitination, or degradation. This approach overcomes a long-standing limitation of ERAD assays, which primarily detect downstream events, and enables direct dissection of the earliest steps that precede ubiquitination, p97-driven extraction, and proteasomal degradation.

Expansion microscopy reveals that the ERQC comprises a dense, multilayered ER architecture that is not resolvable by conventional confocal imaging. Within this compartment, retrotranslocating substrates are selectively enriched and colocalize with the ERAD components HRD1, HERP and OS-9. Furthermore, the cytosolic E3 ligase SCF^FBS2^ also colocalized strongly and selectively with retrotranslocating molecules at the ERQC. These observations support a model in which the ERQC represents a dynamic, stress-responsive reorganization of ER membranes that concentrates ERAD substrates and machinery to facilitate efficient initiation of retrotranslocation and coordination with downstream processing. Our findings extend earlier reports of ERAD substrate accumulation in the ERQC^19–25^ and directly link ERQC architecture to functional retrotranslocation events in mammalian cells.

Collectively, our findings redefine ERAD initiation as a membrane-proximal and reversible sampling process that precedes p97-driven extraction and ubiquitin-dependent proteasomal commitment, and that is spatially organized within the ERQC.

A central outcome of this work is the experimental separation of substrate exposure from extraction. BirA-mediated labeling, protease protection, and subcellular fractionation consistently demonstrate that H2a-BAP becomes transiently accessible to the cytosol while remaining glycosylated and membrane-associated. Deglycosylation coincides with rapid commitment to proteasomal degradation, as deglycosylated intermediates that are no longer membrane-associated accumulate only when proteasomes are inhibited. These observations argue against the existence of a long-lived, freely diffusing cytosolic pool of retrotranslocated substrate. Instead, our findings support a sequence in which substrates undergo initial cytosolic exposure while remaining membrane-integrated, followed by generation of a deglycosylated species that is membrane-proximal but no longer integrated. We also find a substantial membrane-tethered proteasome pool. Together, these data provide direct evidence for tight spatial and kinetic coupling of retrotranslocation, deglycosylation and proteasomal turnover at the cytosolic face of the ER/ERQC (Fig. 7).

**Figure 7.**
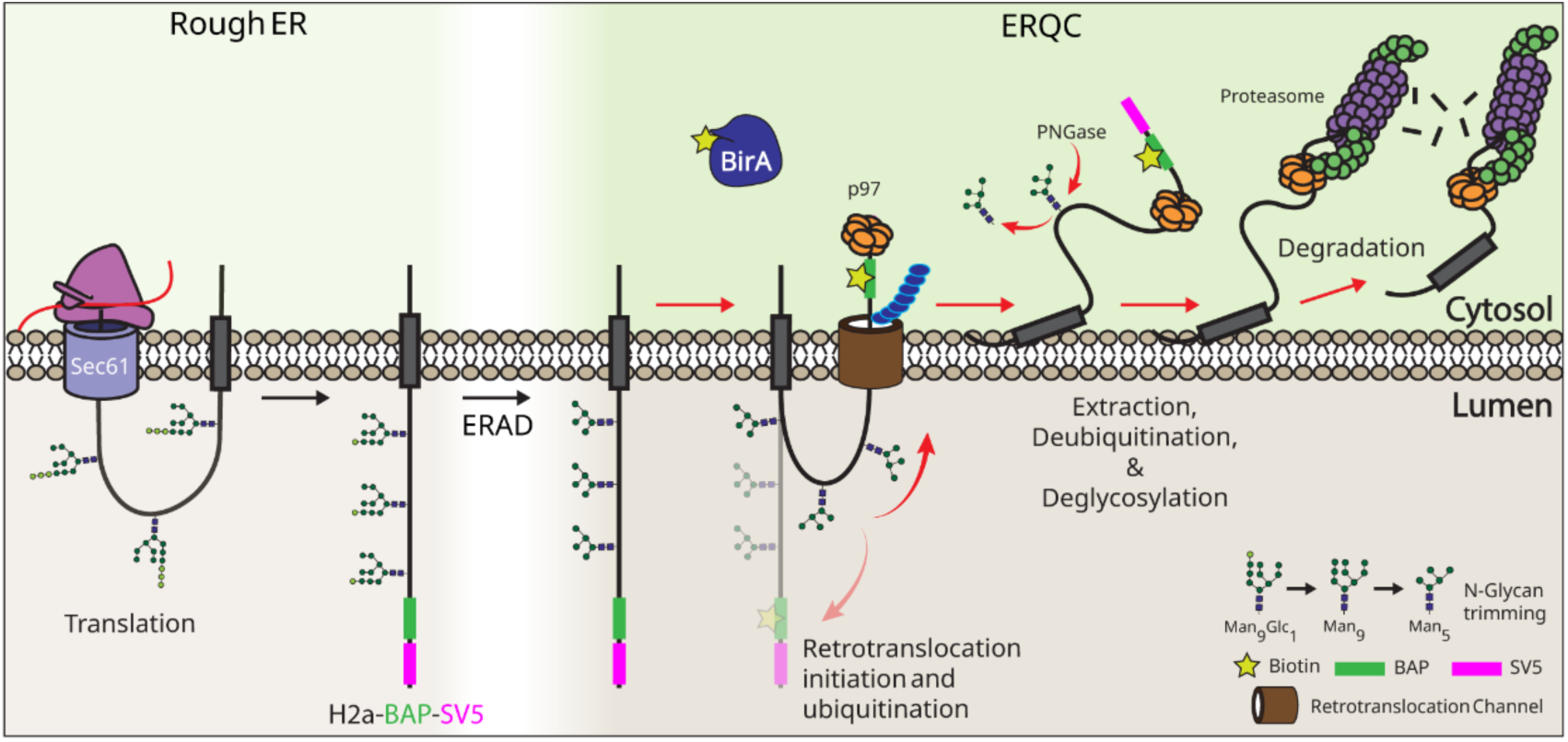
A stepwise model for ERAD initiation at the ERQC. Newly synthesized H2a–BAP–SV5 is co-translationally inserted into the ER membrane via the Sec61 translocon at the rough ER. Its luminal domain is N-glycosylated and after glycan trimming the substrate is routed to the ER-derived quality control compartment (ERQC) for ER-associated degradation (ERAD). At the ERQC, engagement of a retrotranslocation channel initiates retrotranslocation, transiently exposing the luminal tail to the cytosol, thereby permitting BirA-mediated biotinylation of the BAP tag and ubiquitination. In the absence of ubiquitination, the exposed tail can regress back into the ER lumen. Productive ubiquitination commits the substrate to p97-driven extraction from the membrane. During extraction, the transmembrane segment is pulled out, enabling cytosolic deglycosylation by PNGase, followed by ubiquitin chain removal and proteasomal degradation.

Inhibition of the AAA ATPase p97, which is known to extract membrane proteins during ERAD, blocks deglycosylation and degradation but does not prevent initial cytosolic exposure. These results establish that retrotranslocation initiation precedes, and is mechanistically distinct from, p97-dependent extraction and proteasomal degradation, steps that have often been conflated in ERAD models. Importantly, this distinction allows early ER exit events to be interrogated independently of downstream processing and reveals that cytosolic exposure alone is insufficient to commit substrates to degradation.

Protease protection experiments revealed that a substantial fraction of biotinylated H2a-BAP remains luminally protected, largely exceeding what can be attributed to microsome inversion. This observation implies that retrotranslocation involves transient cytosolic exposure followed by partial regression, or back-sliding, prior to irreversible commitment. Cytosol-exposed species are rapidly ubiquitinated and degraded, with polyubiquitinated intermediates accumulating only upon proteasome inhibition. These findings are most readily reconciled with a channel-mediated retrotranslocation mechanism that allows bidirectional substrate movement until stabilized on the cytosolic side by polyubiquitination, consistent with ratcheting models previously proposed for ERAD substrates^59, 65^. While our data do not identify the molecular nature of the retrotranslocation conduit, they demonstrate that exposure and extraction are separable events and establish the ERQC as a functional site of ERAD initiation rather than a passive accumulation zone.

Our perturbation experiments further identify N-glycan processing as an upstream regulatory step. Inhibition of glucose trimming does not prevent retrotranslocation or downstream ubiquitination of H2a-BAP, consistent with glycan-independent ERAD pathways operating alongside lectin-mediated recognition. In contrast, inhibition of mannose trimming markedly reduces retrotranslocation efficiency, positioning mannose trimming upstream of productive engagement with the retrotranslocation machinery. Mannose trimming has previously been shown to be required for ERAD substrate targeting to the ERQC and for degradation^66^, and our data extend these findings by demonstrating its role as a gatekeeper that promotes entry into ERAD rather than directly regulating ubiquitin chain assembly.

Although HRD1 has been proposed to function as a retrotranslocation channel^36–38^, our data indicate that its catalytic E3 ubiquitin ligase activity is dispensable for initial cytosolic exposure of H2a-BAP, but is required for downstream ubiquitination and degradation. Dominant-negative HRD1 strongly impairs polyubiquitination and degradation while leaving retrotranslocation largely intact, resulting in substrate trapping at the ERQC. Similarly, perturbation of the cytosolic SCF^FBS2^ ligase^64, 67^, does not limit retrotranslocation, but traps retrotranslocating molecules at the ERQC. Together, these findings support a model in which multiple E3 ligases act downstream of substrate exposure to enforce irreversible commitment to degradation through polyubiquitination, whereas the initiation of retrotranslocation is governed by distinct upstream determinants.

In conclusion, our results support a stepwise model of ERAD in which misfolded secretory proteins are first targeted to the ERQC through mannose-trimming dependent recognition, undergo transient and reversible cytosolic exposure through a membrane-associated retrotranslocation machinery, and are subsequently committed to degradation through E3-mediated ubiquitination, p97-driven extraction, deglycosylation and proteasomal turnover (Fig. 7). By enabling direct visualization of early retrotranslocation events, the H2a-BAP–BirA system offers a general strategy to capture and dissect initiation steps in ERAD.

## Materials and Methods

### Materials

Tween20 (P9416), N-ethylmaleimide (NEM) (E3876, 500mM stock in EtOH), Biotin (B4639), OptiPrep™ Density Gradient Medium (D1556) and other common reagents were from Sigma-Aldrich. Streptavidin-Agarose was from Millipore (S1638), Protein A agarose beads from Repligen (IPA300S), TEMED from TCI (T0147), Bortezomib from Calbiochem (5043140001), cOmplete Protease inhibitor cocktail from Roche (04693116001). Lactacystin (70980), NB-DNJ (21065), CB5083 (19311) and Kifunensine (10009437) from Cayman chemicals. Promix cell labeling mix ([^35^S]Met plus [^35^S] Cys), 1000 Ci/mmol, was from PerkinElmer Life Sciences.

Expansion microscopy reagents: Acrylamide (AA, A4058), N, Nʹ-methylenebisacrylamide (M1533) and Sodium acrylate (SA, 408220) were from Sigma-Aldrich.

### Cell culture and maintenance

HEK 293, NIH 3T3, and U2OS cells were grown in Dulbecco’s modified Eagle’s medium (DMEM) with high Glucose supplied with 10% Fetal bovine serum (FBS), Penicillin, Streptomycin, and 2mM L-Glutamine, all from Sigma-Aldrich, under 5% CO_2_ at 37°C.

### Antibodies

Streptavidin-HRP (Jackson ImmunoResearch-016-030-084; 1:5000-7500; WB), Rabbit-α-SV5 (CellSignalling-D3H8Q; 1:2000 (WB), 1:750 (IF), 1:250 (ExM)), Mouse-α-SV5 (GeneScript-A01724-100; 1:1000 (WB), 1:750 (IF), 1:250 (ExM)), Rabbit-α-SYVN (Hrd1) (CellSignaling-14773; 1:1000 (WB), 1:100 (IF)), Rabbit-α-H2a (against N-terminus of H2a as described in ^48^) Rabbit-α-Calnexin (Abcam-92573; 1:5000 (WB)), Rabbit-α-RPL26 (CellSignaling-5400; 1:1000 (WB)), Mouse-α-eIF2α (MBL-AT6031; 1:1000 (WB)), Rabbit-α-C9 (20S) (Abcam-ab118902; 1:1000 (WB)), Rabbit-α-Calnexin (SigmaAldrich - C4731; 1:1000 (WB), 1:500 (IF)), Rabbit-α-Calreticulin (CellSignaling-D3E6; 1:400 (IF)). Rabbit-α-Cab45 (1:500 (WB)) and Rabbit-α-Sec61β (1:1000 (WB), 1:500(IF)) as described in ^19^. Rabbit-α-GFP (SantaCruz Biotechnology-SC8334; 1:500 (WB)). Rabbit-α-BiP (1:500 (IF); as described in ^22^. Mouse-α-Myc (CellSignalling-9B11; 1:1000 (WB), 1:1000 (IF)), Mouse-α-HA (BioLegend-MMS-101R; 1:1000 (WB)), Mouse-α-Ubiquitin (CellSignalling-P4D1; 1:1000 (WB)), Mouse-α-FLAG (SigmaAldrich-F3165; 1:1000 (WB), 1:200 (IF)), Mouse-α-S-Tag (Novagen-71549-3; 1:500 (IF)). All the fluorescence secondary antibodies were from Abcam: α-Mouse-Alexa594 (ab150116), α-Mouse-Alexs555 (ab150114), α-Rabbit-Alexa488 (ab150077), α-Rabbit-Alexa647 (ab150083) (1:200 (IF), 1:100 (ExM)). StAv-Cy2 (SigmaAldrich 1:200 (IF)), StAv-Cy3 (SigmaAldrich-GEPA43001; 1:300 (IF)), StAv-Atto647N (SigmaAldrich-94149; 1:200 (ExM), 1:400 (IF)). Secondary antibodies for western blotting were from Jackson ImmunoResearch Labs: Goat-α-Mouse IgG-HRP (115-035-166) and Goat-α-Rabbit IgG-HRP (111-035-144) (1:10000 (WB)).

### Plasmids

H2a-Wild Type in pcDNA1.1 ^48^, H2a-BAP-SV5 (BAP-SV5 ^46^ was cloned with Xba1 before that the C-terminus of H2a-G78R in pcDNA3) ^50^, BAP-H2a-His-Myc in pcDNA3 (BAP was cloned at the N-terminal of H2a with Restriction Enzymes: EcoRI and BamHI), H2a-G78R-Myc in pcDNA3 ^19^, BiP-GFP, BiP-BAP-SV5 (BAP-SV5 tags were cloned between BiP and GFP of the BiP-GFP plasmid with Restriction Enzyme: BamHI). GalT-YFP, HERP-FLAG, and OS91/2-S-Tag (previously described ^22^). BirA and sec-BirA ^46^, FLAG-His-p97 pcDNA3.1 (a kind gift from Dr. Ariel Stanhill), BAP-FLAG-His-p97 (BAP was cloned at the N-terminal of p97 plasmid with Restriction Enzymes: XhoI and BamHI). FBS2 wild type (FBS2-Wt-Myc), mutant FBS2 (ΔFBS2-Myc), HRD1-Wild Type (HRD1-Wt), and HRD1-Dominat-Negative (HRD1-DN) a RING finger mutant cloned in pcDNA3.1 Myc-His vector (previously described in ^22, 64^).

### Transfection, lysis, precipitation and immunoblotting

HEK 293 cells were transfected by the calcium phosphate method. For biotinylation assays, BAP-tagged constructs were co-transfected into HEK 293 cells with equal amounts of BirA or sec-BirA plasmid, and biotin (100 µM) was added for 30 min before harvest. Cell lysis in lysis buffer (1% (v/v) Triton X-100, 0.5% (w/v) sodium deoxycholate (NaDOC), protease inhibitor cocktail, in PBS), immunoprecipitation using Protein A agarose-beads, StAv-precipitation, and immunoblotting was performed as described before ^44^. ECL was performed and the blots were imaged in a BioRad XRS imager.

For glycosidase treatment, Endo H and PNGase F (New England Biolabs) were used according to the manufacturer’s instructions. Ten percent of each lysate or streptavidin-precipitated sample was denatured in 0.5% SDS containing buffer at 100 °C for 10 min and incubated with Endo H for 1 h or PNGase F overnight at 37 °C in the appropriate buffers (supplied with the kit) before SDS-PAGE and immunoblotting.

### Pulse-Chase analysis

[^35^S]-cysteine metabolic labeling was performed as before ^56^. Briefly, cells in 100mm dishes were washed twice with cysteine-free medium, labeled for 20 min in cysteine-free medium containing 200 µCi [^35^S]-Cys. For pulse samples, cells were washed 3 times with cold PBS and pellets were frozen or lysed immediately. For chase samples, after the pulse the label was removed, cells were washed twice with pre-warmed complete medium, incubated in complete medium with puromycin (50 µg/mL) for 1 hour, followed by 1 hour without puromycin, and finally 30 minutes with complete medium supplied with biotin (100µM). Cell pellets were lysed and equal amounts precipitated with α-H2a antibody (as described in ^49^) or StAv-agarose overnight at 4 °C. Precipitated proteins were resolved by SDS–PAGE, and gels were treated with 20% (w/v) 2,5-diphenyloxazole (PPO) before exposure to a phosphor screen in autoradiography cassettes. Radioactive signals were detected using a Fujifilm phosphor-imager (as described in ^66^).

### Fluorescence microscopy

NIH 3T3 and U2OS cells were transfected with Lipofectamine 2000 according to the manufacturer’s instructions, using reagent:DNA ratios of 3:1 and 1:1, respectively. Cells were fixed, permeabilized, blocked, and stained as previously described ^56^. StAv-conjugated fluorophores were applied after secondary antibody staining in 2% BSA/PBS (0.1% Tween-20). Samples were washed three times with 0.1% Tween-20/PBS and left in PBS overnight at 4 °C. Coverslips were mounted the next day using mounting medium on glass slides. Image acquisition was performed with either a Leica LSM 510 or Leica SP8 confocal microscope.

### Expansion microscopy

The protocol was adapted from a previous study ^68^. Cells cultured on 12-mm glass coverslips were treated with Bz (2.5 µM, 5 h) and biotin (100 µM, 3 h), then fixed in 2% acrylamide (AA) and 1.4% (v/v) formaldehyde (FA) in 1× PBS at 37 °C for 5 h. Gelation was performed with a chilled monomer solution (19% sodium acrylate, 10% AA, 0.1% N,N′-methylenebisacrylamide (BIS) in PBS) to which 0.5% APS and 0.5% (v/v) TEMED were added immediately before use. Aliquots of 50 µL gel mix were pipetted onto a pre-chilled parafilm-coated metal plate, and coverslips with AA/FA-treated cells were inverted cell-side down onto the gel drops. Gels were polymerized for 5 min on the cold metal plate, followed by 1 h at 37 °C in a humidified dark chamber.

Coverslips carrying gels were incubated in denaturation buffer (DB; 200 mM SDS, 200 mM NaCl, 50 mM Tris-HCl pH 9.0) with shaking until the gels detached. Detached gels in DB (in 1.5-mL tubes) were boiled at 95 °C for 90 min, washed twice in ddH₂O at room temperature (30 min each), then incubated in fresh ddH₂O overnight at 4 °C for full expansion. The next day, gels were equilibrated in 1× PBS twice for 15 min each before staining. Gels were stained in 2% BSA/PBS, 0.1% Tween-20 at 37 °C in a humid chamber with gentle rocking: primary antibodies, secondary antibodies, and StAv-Atto647 were each applied for 3 h with three ≥10-min washes in 0.1% Tween-20/PBS between steps. Stained gels were washed in PBS/0.1% Tween-20, incubated with DAPI (0.4 mg/mL in PBS, 10 min), washed twice with PBS and then with water, and left in ddH₂O overnight at 4 °C for complete expansion.

Image acquisition: Glass-bottom 6-well plates were coated with poly-L-lysine (0.1 mg/mL in water, 10 min, no wash). Expanded gels were cut into pieces and placed cell-side down on the coated coverslips; excess water was removed, a small drop of water was added on top, and a 25-mm coverslip was placed to form a sandwich. Images were acquired on a Leica TCS SP8 confocal microscope using a 63×/1.4 NA oil objective in Lightning mode (average-max resolution, adaptive strategy) with water as the imaging medium. Three-dimensional stacks were collected with 0.12–0.5 µm z-steps and an x–y pixel size of 35 nm.

### Iodixanol density gradients and carbonate extraction

HEK 293 cells expressing H2a-BAP – BirA, either untreated or treated with bortezomib, were washed with PBS and resuspended in homogenization buffer (0.25 M sucrose,10 mM HEPES (pH 7.4), and 30 mM N-ethylmaleimide (NEM) in water). The cells were passed through a 21G needle five times before homogenization in a Dounce homogenizer (low clearance pestle, 30 strokes). The homogenates were centrifuged at 1000× g for 10 min at 4 °C to remove nuclei and cell debris. Post-nuclear supernatants were subjected to iodixanol gradient centrifugation as described previously ^44^. Two iodixanol step gradients were used: for subcellular fractionation, 10%, 16%, 22%, 28% and 34% iodixanol; and for separation of soluble and membrane fractions, 10%, 13%, 16%, 20% and 24% iodixanol. For carbonate extraction experiments, microsomes were divided equally into three tubes: (1) untreated microsomes (M); (2) SDS-lysed microsomes (S), treated with 1% (w/v) SDS, boiled for 2 min at 95 °C, cooled to room temperature and supplemented with 1% (v/v) Triton X-100 and 0.5% (w/v) sodium deoxycholate (NaDOC); and (3) carbonate-treated microsomes (C), adjusted to 100 mM Na₂CO₃ ^69^. All samples (M, S, C) were incubated on ice for 30 min and then loaded onto iodixanol gradients. One-milliliter fractions were collected, precipitated with 10% trichloroacetic acid (TCA), and analyzed by SDS–PAGE and immunoblotting.

### Protease protection assay (PPA)

Cells were homogenized as per iodixanol protocol, microsomes were subjected to three different treatments as previously described ^50^. Briefly, microsomes were boiled with 1% SDS at 100 °C for 5 min, followed by incubation with or without 0.5 mg/mL proteinase K at 4 °C for 30 min. Or microsomes were directly incubated with proteinase K at 4 °C for 30 min without prior boiling with SDS. The reactions were stopped with 12% TCA; precipitates were boiled, separated on SDS-PAGE and subjected to immunoblotting.

### Statistical analysis and software

Fluorescence images were analyzed on ImageJ as follows: Sample drift in z-stacks was corrected by stack registration using StackReg/TurboReg (translation or rigid-body). Colocalization was quantified with Coloc2 within whole-cell ROIs (excluding nuclei), using Costes automatic thresholding and Costes randomization (≥50 iterations; PSF = 3 pixels). ERQC enrichment was expressed as %ERQC signal = 100 × (background-corrected integrated intensity in ERQC ROI / background-corrected integrated intensity in whole-cell ROI), where integrated intensity was calculated as (mean − background) × area. Western blot band intensities were quantified in Image Lab (Bio-Rad). Graphing and statistical analyses were performed in GraphPad Prism 8. Data are presented as mean ± SD., as indicated in the figure legends. For comparisons between two groups, two-tailed unpaired Student’s t-tests were used. For experiments with more than two groups (or with two independent variables), one-way or two-way ANOVA was used as appropriate, followed by multiple-comparisons tests where indicated. Statistical significance was defined as P < 0.05 (*), P < 0.01 (**), P < 0.001 (***), and P <0.0001 (****).

## Acknowledgements

We would like to thank M. Shenkman and D. Neumann for help with constructs and A. Stanhill for a plasmid. Work was supported by grant 2577/20 from the Israel Science Foundation (GZL).

## Author Contributions

Conceptualization: GZL. Investigation: CP, NOS, HS. Methodology: GP, ORB, CP, NOS. Supervision: GZL. Writing: CP, GZL. Review and editing: CP, NOS, HS, ORB, GP, GZL.

## References

1. Arrigo, A.P., Tanaka, K., Goldberg, A.L. & Welch, W.J. Identity of the 19S ‘prosome’ particle with the large multifunctional protease complex of mammalian cells (the proteasome). Nature 331, 192–194 (1988).

2. Nishikawa, S., Brodsky, J.L. & Nakatsukasa, K. Roles of molecular chaperones in endoplasmic reticulum (ER) quality control and ER-associated degradation (ERAD). J Biochem 137, 551–555 (2005).

3. Olzmann, J.A., Kopito, R.R. & Christianson, J.C. The mammalian endoplasmic reticulum-associated degradation system. Cold Spring Harb Perspect Biol 5 (2013).

4. Oikonomou, C. & Hendershot, L.M. Disposing of misfolded ER proteins: A troubled substrate’s way out of the ER. Mol Cell Endocrinol 500, 110630 (2020).

5. Hosoi, T. & Ozawa, K. Endoplasmic reticulum stress in disease: mechanisms and therapeutic opportunities. Clin Sci (Lond) 118, 19–29 (2009).

6. Oakes, S.A. & Papa, F.R. The role of endoplasmic reticulum stress in human pathology. Annu Rev Pathol 10, 173–194 (2015).

7. Wang, M. & Kaufman, R.J. Protein misfolding in the endoplasmic reticulum as a conduit to human disease. Nature 529, 326–335 (2016).

8. Schubert, U. et al. CD4 glycoprotein degradation induced by human immunodeficiency virus type 1 Vpu protein requires the function of proteasomes and the ubiquitin-conjugating pathway. J Virol 72, 2280–2288 (1998).

9. Lord, J.M. & Roberts, L.M. Toxin entry: retrograde transport through the secretory pathway. J Cell Biol 140, 733–736 (1998).

10. Lencer, W.I., Hirst, T.R. & Holmes, R.K. Membrane traffic and the cellular uptake of cholera toxin. Biochim Biophys Acta 1450, 177–190 (1999).

11. Sandvig, K. & van Deurs, B. Entry of ricin and Shiga toxin into cells: molecular mechanisms and medical perspectives. EMBO J 19, 5943–5950 (2000).

12. Boname, J.M. & Stevenson, P.G. MHC class I ubiquitination by a viral PHD/LAP finger protein. Immunity 15, 627–636 (2001).

13. Schmitz, A., Herrgen, H., Winkeler, A. & Herzog, V. Cholera toxin is exported from microsomes by the Sec61p complex. J Cell Biol 148, 1203–1212 (2000).

14. Tsai, B., Ye, Y. & Rapoport, T.A. Retro-translocation of proteins from the endoplasmic reticulum into the cytosol. Nat Rev Mol Cell Biol 3, 246–255 (2002).

15. Wahlman, J. et al. Real-time fluorescence detection of ERAD substrate retrotranslocation in a mammalian in vitro system. Cell 129, 943–955 (2007).

16. Zhong, Y. & Fang, S. Live cell imaging of protein dislocation from the endoplasmic reticulum. J Biol Chem 287, 28057–28066 (2012).

17. Neal, S., Duttke, S.H. & Hampton, R.Y. Assays for protein retrotranslocation in ERAD. Methods Enzymol 619, 1–26 (2019).

18. Bhaduri, S. & Neal, S.E. Assays for studying normal versus suppressive ERAD-associated retrotranslocation pathways in yeast. STAR Protoc 2, 100640 (2021).

19. Kamhi-Nesher, S. et al. A novel quality control compartment derived from the endoplasmic reticulum. Mol Biol Cell 12, 1711–1723 (2001).

20. Kondratyev, M., Avezov, E., Shenkman, M., Groisman, B. & Lederkremer, G.Z. PERK-dependent compartmentalization of ERAD and unfolded protein response machineries during ER stress. Exp Cell Res 313, 3395–3407 (2007).

21. Leitman, J., Ron, E., Ogen-Shtern, N. & Lederkremer, G.Z. Compartmentalization of endoplasmic reticulum quality control and ER-associated degradation factors. DNA Cell Biol 32, 2–7 (2013).

22. Leitman, J. et al. Herp coordinates compartmentalization and recruitment of HRD1 and misfolded proteins for ERAD. Mol Biol Cell 25, 1050–1060 (2014).

23. Benyair, R., Ogen-Shtern, N. & Lederkremer, G.Z. Glycan regulation of ER-associated degradation through compartmentalization. Semin Cell Dev Biol 41, 99–109 (2015).

24. Cremer, T. et al. RNF26 binds perinuclear vimentin filaments to integrate ER and endolysosomal responses to proteotoxic stress. EMBO J 42, e111252 (2023).

25. Albert, S. et al. Direct visualization of degradation microcompartments at the ER membrane. Proc Natl Acad Sci U S A 117, 1069–1080 (2020).

26. Wiertz, E.J. et al. Sec61-mediated transfer of a membrane protein from the endoplasmic reticulum to the proteasome for destruction. Nature 384, 432–438 (1996).

27. Wilkinson, B.M., Tyson, J.R., Reid, P.J. & Stirling, C.J. Distinct domains within yeast Sec61p involved in post-translational translocation and protein dislocation. J Biol Chem 275, 521–529 (2000).

28. Plemper, R.K., Bohmler, S., Bordallo, J., Sommer, T. & Wolf, D.H. Mutant analysis links the translocon and BiP to retrograde protein transport for ER degradation. Nature 388, 891–895 (1997).

29. Chen, Y., Le Caherec, F. & Chuck, S.L. Calnexin and other factors that alter translocation affect the rapid binding of ubiquitin to apoB in the Sec61 complex. J Biol Chem 273, 11887–11894 (1998).

30. McCracken, A.A. & Brodsky, J.L. Assembly of ER-associated protein degradation in vitro: dependence on cytosol, calnexin, and ATP. J Cell Biol 132, 291–298 (1996).

31. Wang, L., Fast, D.G. & Attie, A.D. The enzymatic and non-enzymatic roles of protein-disulfide isomerase in apolipoprotein B secretion. J Biol Chem 272, 27644–27651 (1997).

32. Knop, M., Finger, A., Braun, T., Hellmuth, K. & Wolf, D.H. Der1, a novel protein specifically required for endoplasmic reticulum degradation in yeast. EMBO J 15, 753–763 (1996).

33. Greenblatt, E.J., Olzmann, J.A. & Kopito, R.R. Derlin-1 is a rhomboid pseudoprotease required for the dislocation of mutant alpha-1 antitrypsin from the endoplasmic reticulum. Nat Struct Mol Biol 18, 1147–1152 (2011).

34. Bordallo, J., Plemper, R.K., Finger, A. & Wolf, D.H. Der3p/Hrd1p is required for endoplasmic reticulum-associated degradation of misfolded lumenal and integral membrane proteins. Mol Biol Cell 9, 209–222 (1998).

35. Hampton, R.Y., Gardner, R.G. & Rine, J. Role of 26S proteasome and HRD genes in the degradation of 3-hydroxy-3-methylglutaryl-CoA reductase, an integral endoplasmic reticulum membrane protein. Mol Biol Cell 7, 2029–2044 (1996).

36. Wu, X. et al. Structural basis of ER-associated protein degradation mediated by the Hrd1 ubiquitin ligase complex. Science 368 (2020).

37. Ye, Y., Shibata, Y., Yun, C., Ron, D. & Rapoport, T.A. A membrane protein complex mediates retro-translocation from the ER lumen into the cytosol. Nature 429, 841–847 (2004).

38. Vasic, V. et al. Hrd1 forms the retrotranslocation pore regulated by auto-ubiquitination and binding of misfolded proteins. Nat Cell Biol 22, 274–281 (2020).

39. Rao, B. et al. The cryo-EM structure of the human ERAD retrotranslocation complex. Sci Adv 9, eadi5656 (2023).

40. Neal, S. et al. The Dfm1 Derlin Is Required for ERAD Retrotranslocation of Integral Membrane Proteins. Mol Cell 69, 306–320 e304 (2018).

41. Nejatfard, A. et al. Derlin rhomboid pseudoproteases employ substrate engagement and lipid distortion to enable the retrotranslocation of ERAD membrane substrates. Cell Rep 37, 109840 (2021).

42. Uta Nakayamada, A.F., Saori Kato, Yuka Kamada, Tsukasa Okiyoneda AMFR provides an ERAD bypass mechanism to maintain proteostasis under canonical E3 ligase deficiency. bioRxiv (2025).

43. Wang, L. et al. TMUB1 is an endoplasmic reticulum-resident escortase that promotes the p97-mediated extraction of membrane proteins for degradation. Mol Cell 82, 3453–3467 e3414 (2022).

44. Shenkman, M. et al. Oligosaccharyltransferase Is Involved in Targeting to ER-Associated Degradation. Cells 14 (2025).

45. Cesaratto, F. et al. BiP/GRP78 Mediates ERAD Targeting of Proteins Produced by Membrane-Bound Ribosomes Stalled at the STOP-Codon. J Mol Biol 431, 123–141 (2019).

46. Petris, G., Vecchi, L., Bestagno, M. & Burrone, O.R. Efficient detection of proteins retro-translocated from the ER to the cytosol by in vivo biotinylation. PLoS One 6, e23712 (2011).

47. Sasset, L., Petris, G., Cesaratto, F. & Burrone, O.R. The VCP/p97 and YOD1 Proteins Have Different Substrate-dependent Activities in Endoplasmic Reticulum-associated Degradation (ERAD). J Biol Chem 290, 28175–28188 (2015).

48. Tolchinsky, S., Yuk, M.H., Ayalon, M., Lodish, H.F. & Lederkremer, G.Z. Membrane-bound versus secreted forms of human asialoglycoprotein receptor subunits. Role of a juxtamembrane pentapeptide. J Biol Chem 271, 14496–14503 (1996).

49. Benyair, R., Ron, E. & Lederkremer, G.Z. Protein quality control, retention, and degradation at the endoplasmic reticulum. Int Rev Cell Mol Biol 292, 197–280 (2011).

50. Sharma, N. et al. The Sigma-1 receptor is an ER-localized type II membrane protein. J Biol Chem 297, 101299 (2021).

51. Spiliotis, E.T., Pentcheva, T. & Edidin, M. Probing for membrane domains in the endoplasmic reticulum: retention and degradation of unassembled MHC class I molecules. Mol Biol Cell 13, 1566–1581 (2002).

52. Kikkert, M. et al. Human HRD1 is an E3 ubiquitin ligase involved in degradation of proteins from the endoplasmic reticulum. J Biol Chem 279, 3525–3534 (2004).

53. Neal, S., Syau, D., Nejatfard, A., Nadeau, S. & Hampton, R.Y. HRD Complex Self-Remodeling Enables a Novel Route of Membrane Protein Retrotranslocation. iScience 23, 101493 (2020).

54. Peterson, B.G. et al. Deep mutational scanning highlights a role for cytosolic regions in Hrd1 function. Cell Rep 42, 113451 (2023).

55. Pisa, R. & Rapoport, T.A. Disulfide-crosslink analysis of the ubiquitin ligase Hrd1 complex during endoplasmic reticulum-associated protein degradation. J Biol Chem 298, 102373 (2022).

56. Ogen-Shtern, N. et al. COP I and II dependent trafficking controls ER-associated degradation in mammalian cells. iScience 26, 106232 (2023).

57. Duriez, M., Rossignol, J.M. & Sitterlin, D. The hepatitis B virus precore protein is retrotransported from endoplasmic reticulum (ER) to cytosol through the ER-associated degradation pathway. J Biol Chem 283, 32352–32360 (2008).

58. Shim, S.M. et al. The endoplasmic reticulum-residing chaperone BiP is short-lived and metabolized through N-terminal arginylation. Sci Signal 11 (2018).

59. Baldridge, R.D. & Rapoport, T.A. Autoubiquitination of the Hrd1 Ligase Triggers Protein Retrotranslocation in ERAD. Cell 166, 394–407 (2016).

60. Benyair, R. et al. Mammalian ER mannosidase I resides in quality control vesicles, where it encounters its glycoprotein substrates. Mol Biol Cell 26, 172–184 (2015).

61. Hurtley, S.M., Bole, D.G., Hoover-Litty, H., Helenius, A. & Copeland, C.S. Interactions of misfolded influenza virus hemagglutinin with binding protein (BiP). J Cell Biol 108, 2117–2126 (1989).

62. Flierman, D., Ye, Y., Dai, M., Chau, V. & Rapoport, T.A. Polyubiquitin serves as a recognition signal, rather than a ratcheting molecule, during retrotranslocation of proteins across the endoplasmic reticulum membrane. J Biol Chem 278, 34774–34782 (2003).

63. Frenkel, Z., Gregory, W., Kornfeld, S. & Lederkremer, G.Z. Endoplasmic reticulum-associated degradation of mammalian glycoproteins involves sugar chain trimming to Man6-5GlcNAc2. J Biol Chem 278, 34119–34124 (2003).

64. Groisman, B., Edward Avezov, and Gerardo Z. Lederkremer The E3 Ubiquitin Ligases HRD1 and SCFFbs2 Recognize the Protein Moiety and Sugar Chains, Respectively, of an ER-Associated Degradation Substrate. Israel journal of chemistry (2006).

65. Hampton, R.Y. & Rine, J. Regulated degradation of HMG-CoA reductase, an integral membrane protein of the endoplasmic reticulum, in yeast. J Cell Biol 125, 299–312 (1994).

66. Groisman, B., Shenkman, M., Ron, E. & Lederkremer, G.Z. Mannose trimming is required for delivery of a glycoprotein from EDEM1 to XTP3-B and to late endoplasmic reticulum-associated degradation steps. J Biol Chem 286, 1292–1300 (2011).

67. Yoshida, Y. et al. Fbs2 is a new member of the E3 ubiquitin ligase family that recognizes sugar chains. J Biol Chem 278, 43877–43884 (2003).

68. Hinterndorfer, K. et al. Ultrastructure expansion microscopy reveals the cellular architecture of budding and fission yeast. J Cell Sci 135 (2022).

69. Fujiki, Y., Hubbard, A.L., Fowler, S. & Lazarow, P.B. Isolation of intracellular membranes by means of sodium carbonate treatment: application to endoplasmic reticulum. J Cell Biol 93, 97–102 (1982).

