## Supplementary material for "ER-associated protein degradation initiates by retrotranslocation from the ER quality control compartment": Suppl.

### Supplementary information

### Supplementary figures

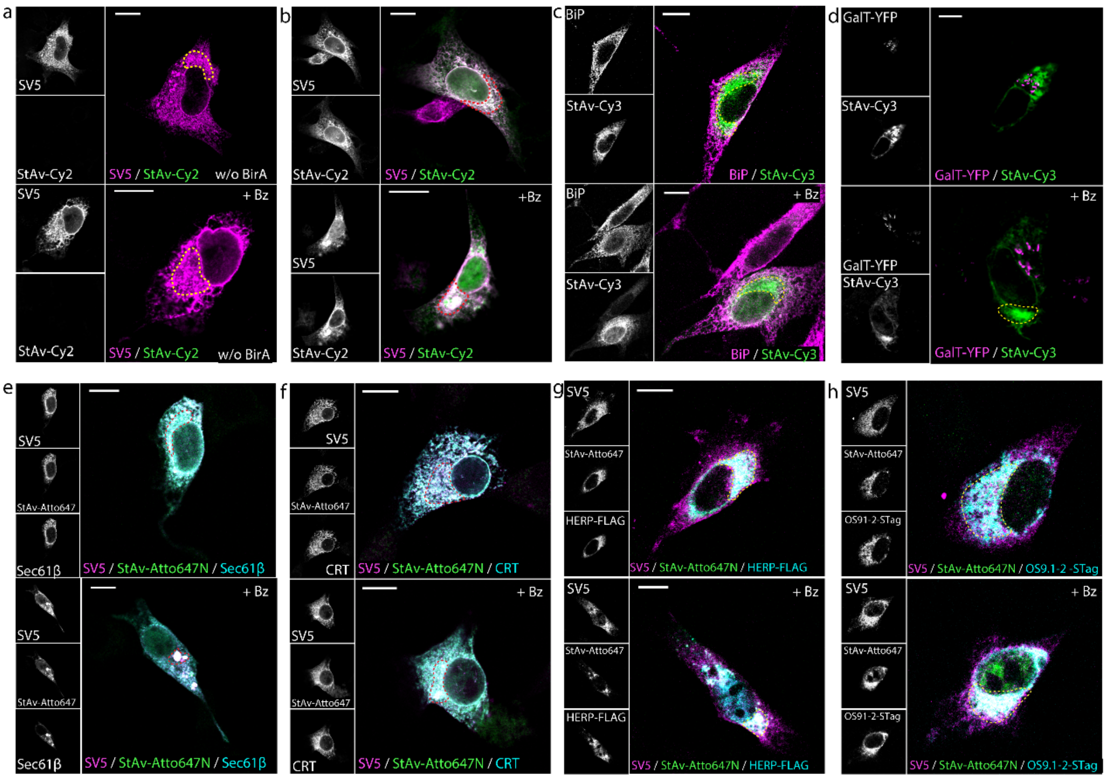


**Supplementary figure 1. BAP-BirA system allows for imaging of retrotranslocation sites in NIH 3T3 cells**: **a.** Immunofluorescence of NIH 3T3 cells expressing H2a-BAP without BirA under basal conditions or after Bz treatment (2.5 µM, 3 hours), stained with α-SV5 and StAv-Cy2 to visualize total H2a-BAP and background biotin signal. **b.** Colocalization of total and biotinylated H2a-BAP in NIH 3T3 cells co-expressing H2a-BAP and BirA under basal and Bz-treated conditions. **c, d.** Colocalization of biotinylated H2a-BAP with BiP (α-BiP) and GalT–YFP (quantified with Coloc2 in Fig. 1o). **e, f.** Colocalization of total H2a-BAP (α-SV5) with biotinylated

H2a-BAP and ER markers Sec61β and CRT. **g, h.** Colocalization of total H2a-BAP with biotinylated H2a-BAP and ERQC markers HERP-FLAG (α-FLAG) and OS9.1/2–S-tag (α-S-tag). Yellow/red dashed ROIs mark the ERQC.  Scale bars= 10 μm.

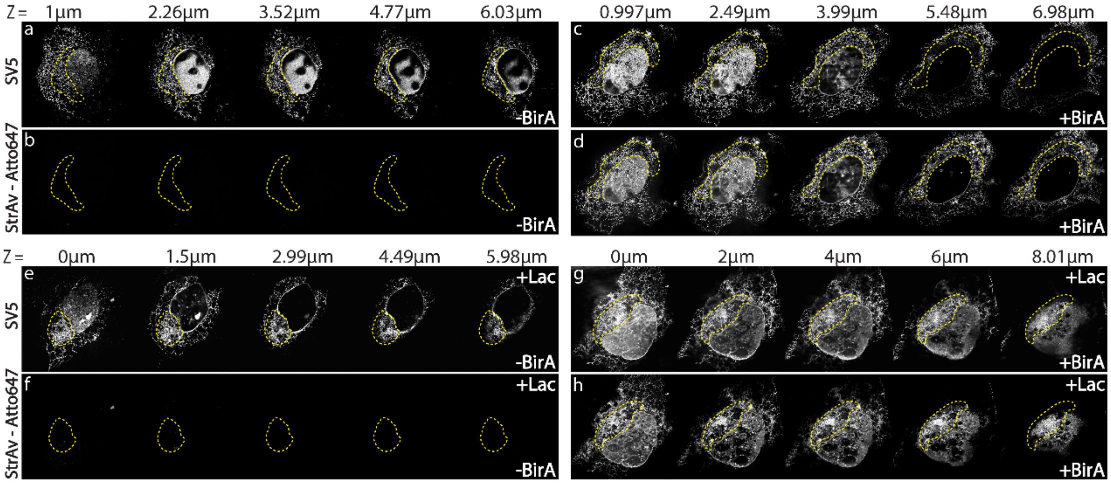


**Supplementary figure 2. High-resolution imaging of the ERQC by expansion microscopy:** U2OS cells expressing the H2a-BAP – BirA system (Fig. 2) were processed for expansion microscopy; montages show five optical sections along the z-axis for each channel in cells expressing H2a-BAP alone under basal conditions (a, b), H2a-BAP together with BirA under basal conditions (c, d), or H2a-BAP–BirA after bortezomib treatment (Bz, 2.5 μM, 3 hours; e-h). The distance from the bottom of the cell is indicated for each z-section and the ERQC is outlined with yellow dashed lines.

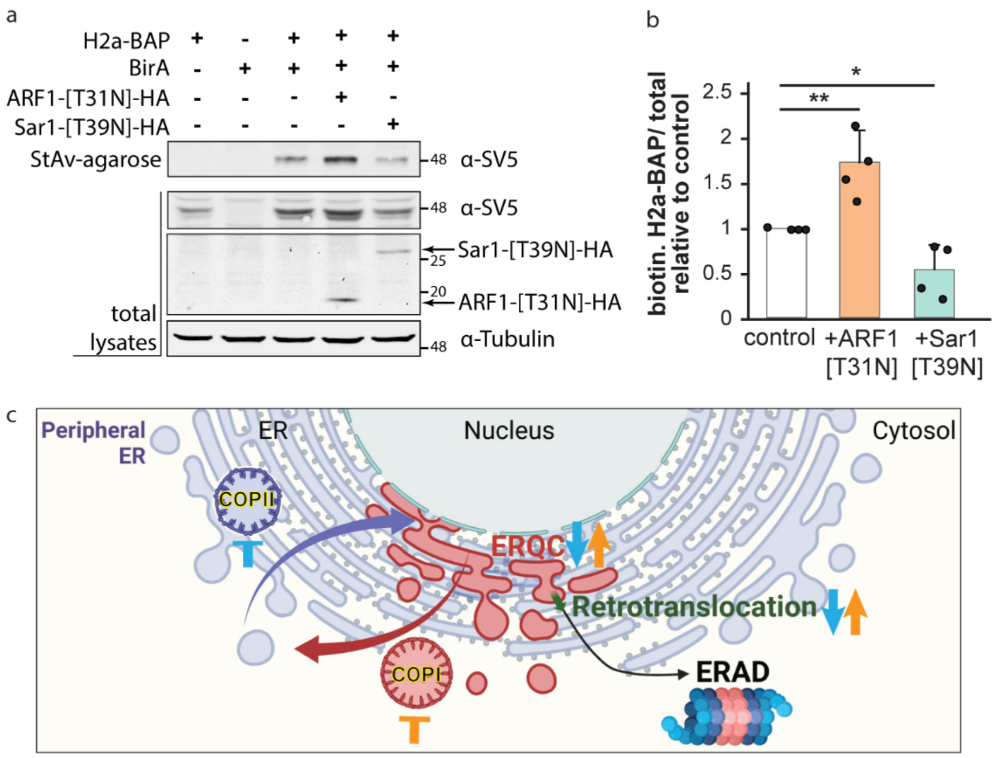


**Supplementary Figure 3. Perturbing COPI/COPII trafficking modulates H2a-BAP retrotranslocation: a.** HEK 293 cells expressing H2a-BAP - BirA together with ARF1-T31N-HA or Sar1-T39N-HA were lysed; 10% of the lysate was analyzed by western blot for total H2a-BAP (α-SV5), α-HA and α-tubulin, and the remaining lysate was subjected to StAv-agarose precipitation and probed with α-SV5 to detect biotinylated H2a-BAP. **b.** Biotinylated H2a-BAP signals (from a) were normalized to total H2a-BAP and expressed relative to control (mean of 4 experiments ± SD; unpaired t-test with Welch’s correction). **c.** Scheme illustrating COPI- and COPII-mediated vesicular transport of ERAD substrates between the peripheral ER and the ERQC (the scheme was designed on BioRender).

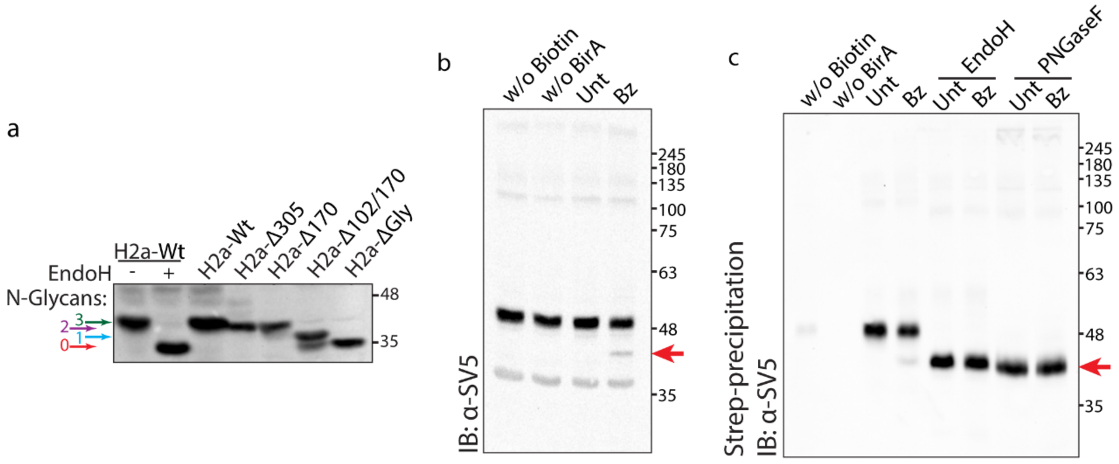

**Supplementary figure 4. Proteasome inhibition causes accumulation of soluble deglycosylated H2a-BAP. a.** H2a-Wt, expressed in HEK 293 cells, was immunoprecipitated and treated or not with Endo H, and compared by Western blot with H2a-Wt and H2a glycosylation mutants (H2a-Δ102, H2a-Δ305, H2a-Δ102/170 and H2a-ΔGly (lacking all 3 N-glycosylation sites)); colored arrows indicate the number of N-glycans. **b.** HEK 293 cells expressing the H2a-BAP - BirA system were left without biotin, without BirA, or were treated with bortezomib (Bz, 2.5 μM, 3 hours) as indicated, and analyzed on western blot to detect total H2a-BAP (α-SV5). **c.** Lysates (from a) were precipitated with StAv-agarose and were mock treated or incubated with Endo H or PNGase F to deglycosylate biotinylated H2a-BAP, revealed by α-SV5 (b). The deglycosylated species is indicated by the red arrow.

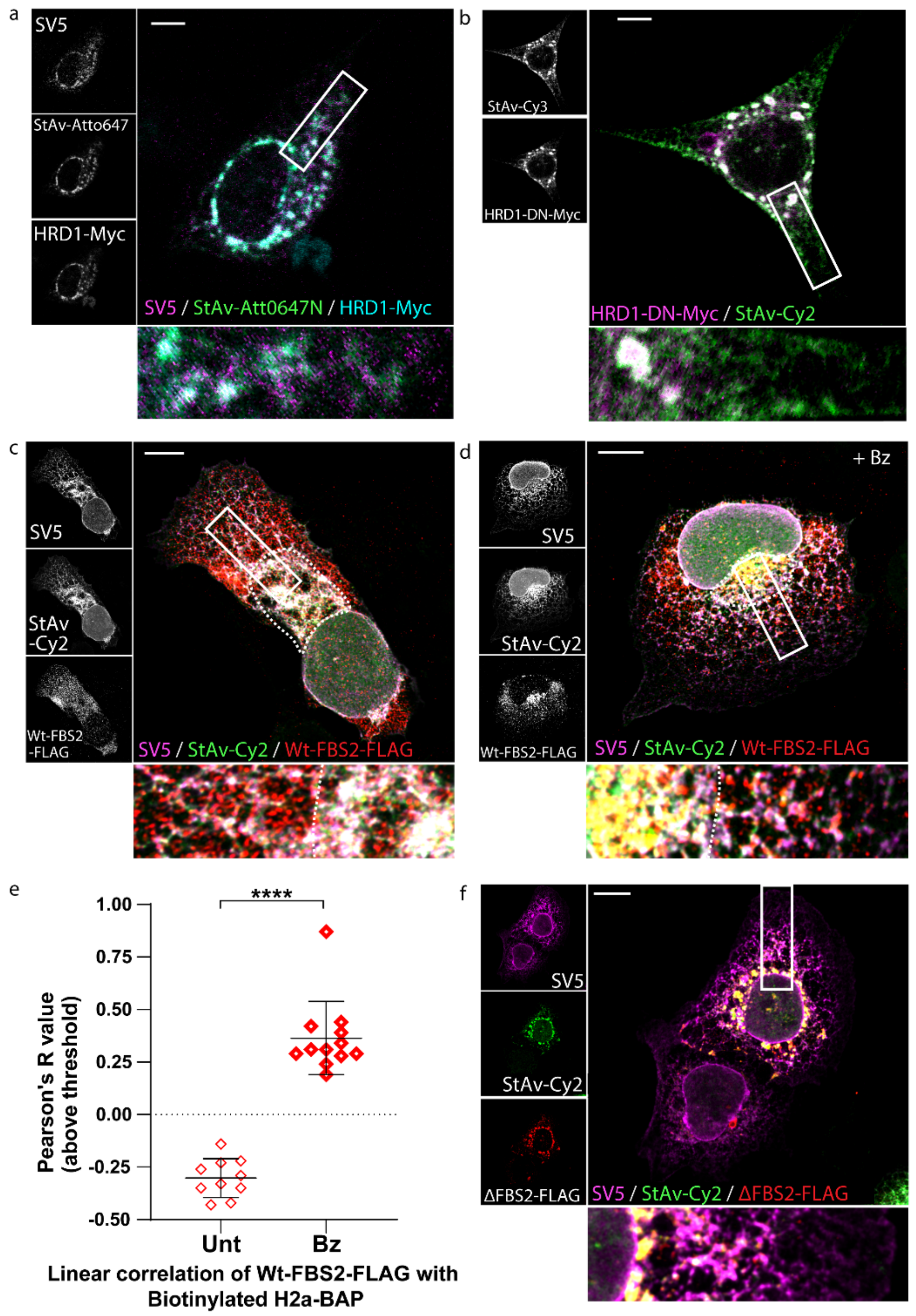


**Supplementary Figure 5. Dominant negative mutants of membrane and cytosolic E3 ligases trap retrotranslocating H2a-BAP at the ERQC: a, b.** NIH 3T3 cells expressing the H2a-BAP–BirA system together with WT HRD1-Myc (a) or dominant-negative HRD1-DN-Myc (b) were fixed and stained for total H2a-BAP (α-SV5), biotinylated H2a-BAP (StAv-Atto647 or StAv-Cy2) and HRD1/HRD1-DN (α-Myc); boxed regions are shown enlarged below. **c, d.** U2OS cells expressing the H2a-BAP - BirA system together with Wt-FBS2-FLAG, alone or with Bz, were stained for total H2a-BAP (α-SV5), biotinylated H2a-BAP (StAv-Cy2) and FBS2-FLAG, and imaged by confocal microscopy. **e.** Pearson's R value calculated above the threshold using

Coloc2 of the whole cell (from c, d). **f.** U2OS cells expressing the H2a-BAP - BirA system together with ΔFBS2-FLAG to assess the impact of FBS2 and its F-box deletion on ERQC accumulation of retrotranslocating H2a-BAP (highlighted with dashed lines). Scale bars= 10 µm.
